# A Characterisation of the Invasive Lema Beetle, *Lema equestris* (Coleoptera: Chrysomelidae), in Hawai‘i

**DOI:** 10.64898/2026.04.28.721477

**Authors:** Mikinley Weaver

## Abstract

Introduced to Hawai‘i in 2019, *Lema equestris* has become a garden pest commonly reported on *Solanum americanum,* which is grown as a native and cultural plant in Hawai‘i and supports native vertebrates elsewhere across Oceania. Originally identified as *L. solani,* the species was later found to have been misidentified. Here, molecular and morphological evidence is used to discriminate Hawaiian specimens from *L. solani* and support the updated identification of *L. equestris*. As a new invasive species, it is important to confirm host associations and determine whether it will prey on important species, such as endemic or endangered plants, in its new range or any potential range to which it could spread. To this end, feeding assays were performed with adults, first-instar larvae, and newly hatched naïve larvae on 11 potential hosts, comprising mostly *Solanum* species: *Solanum americanum,* potato, tomato, tomatillo, poha (gooseberry), chili pepper, eggplant, tobacco, tree tobacco, cabbage, and Brazilian nightshade. While feeding was attempted on cabbage, poha berry, and Brazilian nightshade, no host besides *S. americanum* supported survival. Rearing was used to further characterize the biology and life history of *L. equestris*, including instar length and distinctive morphological traits for identifying each life stage. While many basic biological traits are confirmed here, much remains to be studied to better understand this species and why it has begun to spread.

## INTRODUCTION

Oceania holds 16.7% of the world’s biodiversity hotspots, with the Pacific islands comprising half of that biodiversity (Taylor & Kumar, 2016), but this biodiversity is under constant threat (Jupiter *et al*., 2014; Keppel *et al*., 2014). The Pacific islands alone claim 150 critically endangered terrestrial vertebrates across 23 countries (Kumar & Tehrany, 2017), 50% of Pitcairn fauna are threatened (Waldren *et al*., 1995), 2% of vascular plants are extinct and 32% threatened in Niue (Heenan *et al*., 2026), and at least 15% of plants are threatened in Palau with 61% too data deficient for a determination (Costion *et al*., 2009). Serving as a sentinel of the biodiversity challenges in Oceania, Hawai‘i is home to over 9,975 endemic species (Evenhuis & Miller, 2015), with many imperilled or extinct, earning the islands the nickname the “extinction capital of the world.” Across the islands, 10% of native plants have gone extinct since 1840 (Wood *et al*., 2019), and 37.9% of native Lepidoptera have gone extinct within the last 100 years (Austin & Rubinoff, 2024). On top of this, many more species are threatened or endangered, with some having only a few individuals remaining in the wild, and some existing only in captivity (Rønsted *et al*., 2022). While the specific reasons for these extinctions vary, most result from anthropogenic pressure, including the introduction of non-native species. Introduced species can outcompete natives (Cox & Elmqvist, 2000; Meyer, 2004; Feenstra & Clements, 2008; Baret *et al*., 2013), precipitate habitat destruction (DeNitto *et al*., 2015; Peck *et al*., 2026), or directly target native species (Kleeck & Holland, 2018). Because, like many island nations, Hawai‘i is economically dependent on travel, there is a heightened risk of foreign species being introduced, with an average of 22 taxa introduced and 4 established each year (Reichard & White, 2001). Despite extensive interagency efforts to develop a comprehensive biosecurity program (Arnott *et al*., 2021), additional invasive species continue to be established (Burbano *et al*., 2011; Paudel *et al*., 2023; Villalobos *et al*., 2024).

These introduced species can threaten native plants that are endangered or culturally important, such as those in the genus *Solanum.* There are five *Solanum* species native to Hawai‘i, with the presence of a sixth debated. The first four are endemic and considered endangered or critically endangered (USFWS, 2019; McClelland *et al*., 2020; IUCN, 2024) and include pōpolo kū mai (*Solanum incompletum,* Dunal), pōpolo (*S. nelsonii,* Dunal), Nihoa pōpolo (*S. caumii,* D.McClelland), and pōpolo aiakeakua (*S. sandwicense,* Hook.) (Münster & Wieczorek, 2007; McClelland *et al*., 2020). The fifth species is the common *S. americanum* Mill, and the debated sixth is *S. opacum* A.Braun & C.D.Bouché. The identification of *S. opacum* is complicated by the difficulty of distinguishing it morphologically from *S. americanum*, but it has been claimed to be present in Hawai‘i (Särkinen *et al*., 2018). For this manuscript, both *S. opacum* and *S. americanum* are treated as *S. americanum,* though differences may be gleaned if proper identifications can be made. Commonly called edible pōpolo, *S. americanum* has been used as an edible and medicinal plant by native Hawaiians (Abbott & Shimazu, 1985; Särkinen *et al*., 2018). It is also sold by some plant nurseries for its indigenous status and cultural uses. However, a recent chrysomelid beetle introduced to Hawai‘i, *Lema equestris* Lacordaire, 1845, poses a threat to this practice.

Originally from Central America (Lacordaire, 1845), *L. equestris* was first found in Hawai‘i beginning in 2019, where it was erroneously identified as *L. solani* Fabricius, 1798. Since its introduction, *L. equestris* has become a commonly reported garden pest, feeding on edible pōpolo and, anecdotally, damaging *Physalis peruviana* L. Because no host or life history has been reported for *L. equestris*, assumptions about host preferences and pest tendencies have been based on the morphologically similar *L. solani*, which has undergone recent range expansions through international trade, thereby becoming a pest species. In its native range, *L. solani* is found across the Eastern and Southern United States from Delaware to Texas, has been found in Taiwan since 2013 (Lee & Matsumura, 2013), and one individual was sighted in Hong Kong in 2024, but it is likely not established. In its native range, *L. solani* is reported from *S. americanum,* but records exist for potato [*S. tuberosum* L.], *S. carolinense* L., and tobacco [*Nicotiana tabacum* L.] (Clark, 2004), as well as *S. nigrum* L., an unknown type of bean, “*Bombeya*” [possibly a misprint of the genus *Dombeya* according to Clark (2004)], and cabbage [*Brassica oleracea* L.] (White, 1993). In total, *L. solani* is reported to utilize six solanaceous hosts, one Malvaceae host, one Fabaceae host, and one Brassicaceae host. Another similar beetle, *L. balteata* JL LeConte, 1884, which has been suggested to be synonymous with *L. equestris* despite having a different range, is reported to feed on *Physalis* and *S. americanum* (Clark, 2004).

As these beetles are becoming increasingly relevant to Hawaiian residents seeking to protect their property and native species, it is important to confirm the species present on the islands and their relevant life-history characteristics. If the diversity of hosts on which *L. solani* can feed is representative of *L. equestris*, it would indicate a latent agricultural threat that could soon manifest and may spread to other islands or countries within Oceania that have sufficient populations of host plants. Host specificity of Hawaiian specimens of *L. equestris* was assessed with host-choice tests, and multiple generations of the beetles were reared to describe the species’ biology and lay the foundation for further studies on how this biology can be leveraged to control the species outside its native range.

## MATERIALS AND METHODS

### Molecular Identification

DNA was extracted from six wild-caught *Lema equestris* beetles, two larvae (one L2, one L3), and four adults (two males and two females) and molecular identification was utilized to confirm species identity. One male and one larva were collected from Carlsmith Beach Park (19°44’00.9“N 155°01’40.9“W), while the other specimens were collected from the University of Hawai‘i Hilo (19°42’00.3“N 155°04’58.2“W). DNA extractions were performed as part of a microbiome project (Mason *et al*., 2026) using the Zymobiomics Magbead 96 Kit (Zymo Research, Tustin, CA, USA). Samples were extracted with the lysis step utilizing a Geno/Grinder 2010 (SPEX SamplePrep Metuchen, NJ, USA) for five rounds of 5 min at 1,750 RPM with 5-min rest, and the extraction automated with a Kingfisher Apex (Thermo Fisher Scientific Inc., Waltham, MA, USA) following the manufacturer’s protocols. For comparison, two *Lema solani* specimens were collected in November 2025 from Secret Lake Park in Orlando, Florida (28°40’30.7“N 81°19’35.6“W). One was preserved in DNA/RNA shield (Zymo Research) and frozen for DNA extraction, while the other had one leg removed for DNA extraction, which was placed in DNA/RNA shield and then pinned for morphological comparison. Lysis of *L. solani* specimens was performed using 0.2 mm glass beads in microcentrifuge tubes, shaken for 15 mins with a Disruptor Genie (USA Scientific Inc., Ocala, FL, USA), and the lysate was manually extracted using a Zymo Magbead Kit. Purified DNA was amplified using Q5 Hotstart high-fidelity polymerase 2x master mix (New England Biolabs, Ipswitch, MA, USA) and primers at a final concentration of 0.5 nM. The conditions were as follows: 98C for 30s; 31 cycles of 98C for 30s, 55C for 30s, 72C for 2 min; and final extension of 72C for 10 min. Amplicons were generated for 18S, 28S, and COI mitochondrial barcoding genes, purified using the DNA Clean & Concentrator MagBead Kit (Zymo Research), and Sanger sequenced.

A consensus sequence was created using Geneious Prime (https://www.geneious.com) and compared with publicly available NCBI Genbank sequences (accession numbers listed in Supplemental Table S1). Maximum-likelihood trees were constructed using W-IQ-Tree (Trifinopoulos *et al*., 2016). ModelFinder (Kalyaanamoorthy *et al*., 2017) was used to select the best model, resulting in using the TIM2+F+I+G4 model with bootstrap branch support performed with UFBoot2 at 1000 replicates (Hoang *et al*., 2018). Tree branches were coloured according to species with the RainbowTree tool (Paradis, 2012) for the 19 most relevant species. The 18S, 28S, and COI genes were compared between *L. solani* and *L. equestris* as single-gene comparisons, while a combination of 18S and COI was used for the multigene analysis.

### Morphological Examination

Ten individuals of *L. equestris* were collected from three sites on O‘ahu and six on Hawai‘i island (totalling 90 individuals): One‘ula Beach Park (21°18’22.5“N 158°01’43.5“W), the Oceanic Institute (21°18’53.8“N 157°39’58.5“W), Kaneohe Stream/Kamehameha Hwy (21°24’43.8“N 157°47’58.8“W), Waiakahi‘ula Beach Park (19°33’38.7“N 154°53’19.6“W), Carlsmith Beach Park, Onekahakaha Beach Park (19°44’15.3“N 155°02’22.2“W), Naniloa Golf Course (19°43’40.5“N 155°03’54.5“W), USDA-ARS Pacific Basin Agricultural Research Center (19°41’51.9“N 155°05’38.9“W), and the University of Hawai‘i Hilo. Specimens were examined for morphological differences, and then four from each location (2 males and 2 females) were dissected to observe internal anatomy. Two specimens of *L. solani*, as previously described, were collected from Orlando, Florida, USA, and examined while alive before one was used for DNA examination. In addition to physical specimens, iNaturalist was explored for a total of 82 records of Hawaiian *L. equestris* (labeled as *L. solani*), 350 records of *L. solani* in their native range, 18 records of *L. balteata*, and 47 records of *L. equestris* in their native range. iNaturalist records were filtered to include only adults, and misidentifications were discarded.

The type specimen for *L. balteata* is available as a series of high-resolution images from the Museum of Comparative Zoology (MCZ:Ent:4257), but type specimens for *L. solani* and *L. equestris* could not be located for comparison. All the examined specimens (physical and those found on iNaturalist) were compared and sorted into phenotypes based on shared characteristics. These phenotypes were then compared with the original species descriptions and literature to determine the species identity.

### Rearing and Life Stages

The colony was started by collecting leaves with eggs from populations collected from both the Oceanic Institute, Waimanalo, and Onekahakaha Beach Park, Hilo Hawai‘i, USA. The leaves were placed in individual plastic cages (4 cm x 4 cm) and kept in the laboratory at 25±2°C until the larvae emerged. First-instar larvae were separated into individual petri dishes containing a small piece of filter paper moistened daily and a single leaf of *S. americanum,* which was replaced as it was consumed. The petioles of the leaves were placed in microcentrifuge tubes with holes in the lids, which were filled with distilled water. To track life-stage durations, the larvae were examined at least twice daily and checked for exuviae or noticeable changes in head capsule size. This allowed each instar duration to be identified with an accuracy of 12 hours or less. The growth rates of three replicates of five larvae (n=15) were tracked by photographing them across their larval period and measuring them using ImageJ (Fiji 2.16.0; Schindelin *et al*., 2012). Once larvae reached the prepupal stage, the leaves were removed, and the dish was placed in a dark box to encourage pupation, allowing pupae to be observed inside their pupal cases through the clear dish. Adults post-eclosion were placed in a rearing cage containing a potted *S. americanum* plant and were allowed to breed, mate, and oviposit freely. The colony was allowed to go through two generations before being restarted, and this process occurred four times. Once with the Waimanalo population in March 2023, and three times with the Hilo populations in June 2023, September 2024, and December 2024.

### Diet Preference and Host Plant Trials

#### I. Adult no-choice feeding assay

No-choice tests were performed at least three times per plant with adult beetles collected in Hilo, HI, to confirm host reports. Each host species was tested individually by haphazardly selecting a minimum of five beetles from the colony and depriving them of food for at least 36 hours. The beetles were then placed into a feeding arena consisting of a loosely covered plastic petri dish (100 mm × 10 mm) with ∼2000 mm^2^ of excised foliage to be tested, with leaf petioles placed in a water tube to prevent humidity fluctuations within the dish. Beetles were observed every hour for the first 12 hours to record their locations in the arena, and whether they were feeding on the host plant. If damage was noted, the feeding scars were imaged and measured with ImageJ to determine the area consumed by comparing with images taken before the assay. The beetles were then left undisturbed for another 12 hours. After 24 hours in the arena, the test leaves were removed for examination and replaced with *S. americanum* leaves for one hour (similar to a B-A assay). During this recovery period, feeding was recorded, but not measured, to demonstrate readiness to feed (Van Driesche & Murray, 2004), and all feeding measurements for *S. americanum* came from independent trials. Removed beetles were placed in a separate cage to ensure that selected beetles were always assay-naïve. The trial diets consisted of leaves of edible pōpolo [*S. americanum*], Brazilian nightshade [*S. seaforthianum* Andr.], potato [*S. tuberosum*], tomato [*S. lycopersicum* L.], eggplant [*S. melongena* L.], poha berry [*P. peruviana*], tomatillo [*P. philadelphica* Lam.], cabbage [*B. oleracea*], cultivated tobacco [*N. tabacum*], and tree tobacco [*N. glauca* Grah.].

#### II. Adult potted plant choice trial

A choice test was conducted with 5 adult beetles, repeated 3 times, released into a 0.25 m^3^ flight cage containing potted edible pōpolo, poha berry, cultivated tobacco, and potato. All plants were young (<40 cm in height) and potted individually. Adult beetles were allowed to roam freely within the cage for 48 hours. After 48 hours, the plants were removed, and any feeding scars or egg masses were identified.

#### III. Larval no-choice feeding assay

No-choice tests were also performed on first-instar larvae within 12 hours post-hatching. This age was chosen because it is a critical growth stage during which larvae feed continuously, except when moving short distances on the host plant, and grow significantly. The setup of the feeding arena was similar to the adult tests except that before the leaves were placed in the arena, they were damaged in two areas by squeezing them with fine-tipped forceps to release volatiles that might elicit feeding. Three replicates of at least five L1 larvae were placed directly on the test leaves for 4 hours and observed hourly for feeding activity. Unlike in the adult trials, larvae did not have food withheld; instead, they were removed directly from the host, *S. americanum*, and placed onto the test leaves (an A-B style assay). Preliminary trials indicated that waiting 4 hours before returning larvae to their original host plant provided sufficient time to observe feeding behaviour. Larval feeding was tested on edible pōpolo, potato, tomato, poha berry, tomatillo, cabbage, eggplant, tree tobacco, and cultivated tobacco. To confirm potential feeding responses, a secondary test was performed using poha berry and cabbage, in which L1 larvae were left on the leaves for 24 hours to observe long-term feeding behaviour.

#### IV. Larval host-naïve no-choice feeding assay

Brazilian nightshade, tomato, chili pepper [*Capsicum annuum* L.], cabbage, and poha berry plants were tested with host-naïve larvae, or L1 larvae that eclosed from eggs placed on the test species rather than being allowed to hatch on *S. americanum* first. Cabbage, Brazilian nightshade, tomato, and poha berry were tested by splitting a clutch of eggs and placing three replicates of at least three unhatched eggs each onto a leaf of the test plant. Leftover eggs from the clutch (≥4) were left to hatch on the host plant (*S. americanum*) as control larvae. After hatching, the larvae were observed four times a day. If any larvae were observed to have left the test plant, they were gently returned to the plant using a soft brush. Larvae were kept on these test leaves for three days to observe their growth. Chili pepper was tested using the same methods, except that, due to a lack of feeding response in other non-typical host trials and limited eggs during the time when chili pepper was available, two replicates of four eggs were used.

### Data Analysis

To evaluate how feeding varied by plant species and life stage, a gamma hurdle model was fitted to the leaf area consumed in each trial. A Bayesian approach was used because 16 of the 24 plant-life stage combinations showed no feeding, which produces separation under maximum-likelihood methods, and partial pooling across combinations stabilises estimates drawn from low replicate numbers.

The model has two components, the hurdle and the gamma model (Supplemental Data Equations S1-S3). The hurdle is the probability that any feeding occurred, as evidenced by feeding damage. The second component, the Gamma model, is for the amount consumed given that the replicate fed, with the mean scaling linearly with the number of beetles and the duration of the trial, so that mu is the mean consumption per beetle present per hour of exposure. Because of the orders of magnitude difference in beetle consumption by life stage, both components used a non-centred multilevel structure where each life stage had its own mean and combinations are partially pooled toward the mean of their own life stage. Three quantities were generated for each combination: probability of feeding, p; the mean feeding rate, mu; and their product p * mu, the unconditional expected consumption rate. Checks were performed to ensure that the model fit was appropriate and are further described in the supplemental (Supplemental Figures 1, 3, and 5).

Posterior draws were used to compute pairwise contrasts between plant species within each life stage, as differences and as ratios (Supplemental Data Equation S7), and posterior predictions at each combination’s own trial design (Supplemental Data Equation S9). To test the null hypothesis that feeding was negligible for non-hosts, regions of practical equivalence (ROPE) were specified before fitting, based on ∼10% of the lowest raw consumption rate per life stage (L1 and adult). The resulting upper bound of the ROPE determined for expected rate (p * mu) was 0.4 mm^2^ beetle^−1^ h^−1^ for adults, 0.02 for first instars and host-naïve larvae (Supplemental Data Equation S8). For differences in p the region chosen was 0.10. The posterior probability that a combination or contrast falls within the relevant region was calculated alongside every estimate (Supplemental Figure 6).

Survival and length were analysed with a second Bayesian model (Supplemental Data Equations S4-S6). Survival was modelled as a staged process with two binomials: larva that survived to 24 h and those that survived to the end of the trial. Length, using final body length as the variable to evaluate stunting, was modelled on the log scale so that the plant-versus-control contrast is a ratio, and is reported as a percentage of control length. A region of practical equivalence of 0.10 was specified before fitting, below which survival is not taken to indicate that a plant supports development. As before, checks were performed to ensure that the model fit was appropriate and are further described in the supplemental (Supplemental Figures 2 and 4).

Data analysis was performed with R v4.6.0 (R Core Team, 2026) with models were fit using cmdstanr (Gabry et al., 2026). Posterior draws were processed using tidybayes (Kay, 2023) and visualised with ggplot2, ggdist (Kay, 2024) and bayesplot (Gabry & Mahr, 2025). Eight chains were run for 4,000 iterations each, discarding the first half as warm-up (16,000 retained draws), with adapt_delta = 0.99, a maximum tree depth of 12 and a fixed seed. Across all parameters and derived quantities of the feeding model the maximum rank-normalised split R-hat was 1.002, the minimum bulk- and tail-effective sample sizes were 5,907 and 8,294, and the minimum E-BFMI was 0.85; for the survival and growth model the corresponding values were 1.002, 4,370, 6,537 and 0.83. Neither model produced a divergent transition or an iteration saturating the maximum tree depth. Rank plots and pairwise scatterplots were inspected for divergences and funnelling. Further details can be found in the supplemental (Supplemental Text 1-1.9 and Figures 1-6).

**Figure 1.**
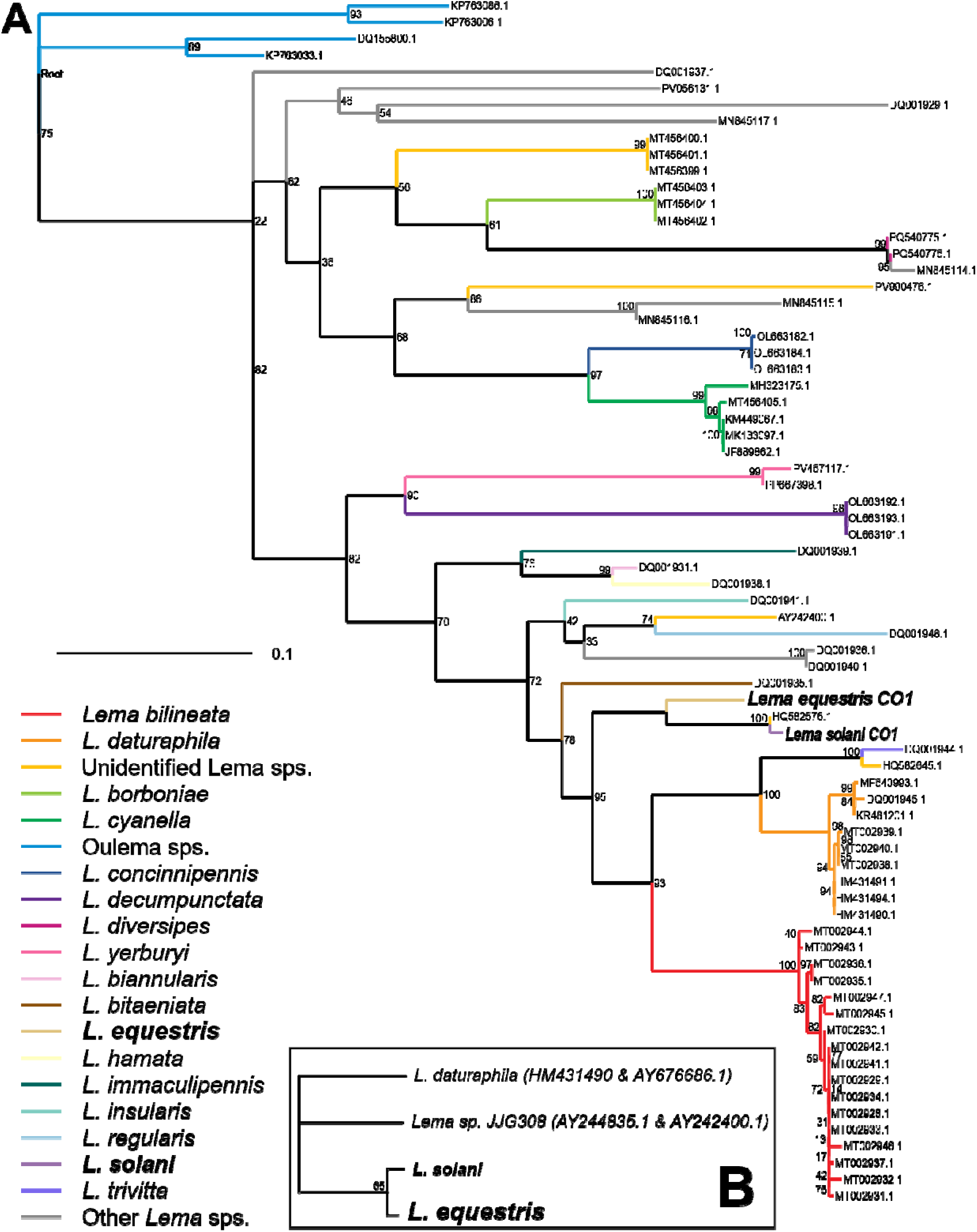
Phylograms of *Lema* species. A. COI sequences with an *Oulema* outgroup. B. Multi-gene analysis combining 18S and COI genes. Numbers at each branch represent the bootstrap support values, with branch lengths representing genetic distance.

## RESULTS

### Molecular Identification

Barcodes were successfully amplified across all tested barcoding regions in *L. equestris* and *L. solani*, and all the samples had identical sequences for each respective species. Pairwise comparisons revealed little variability in the 18S and 28S genes between species, with a greater difference in the more commonly used COI barcoding gene (18S – 99.6%, 28S – 99.7%, COI – 92.6% pairwise identity). Single and multigene analyses found *L. equestris* and *L. solani* to be closely related but supported their distinction as separate species (Figure 1).

### Morphological Examination

Specimens of *L. solani* collected in Florida presented with a distinct colouration, while Hawaiian specimens could be distinguished into two similar but distinct phenotypes. The distinct *L. solani* colouration was a beetle with a solid rufous elytral margin and subapical fascia (Figure 2), while the two Hawaiian specimens phentypically categorized as those with a rufous prothorax and a black elytral margin interrupted with a rufous medial fascia and subapical fascia (Figure 3), or those with a black prothorax, an interrupted elytral margin, and an absent subapical fascia (Supplemental Figure 7C). Each phenotype matched the descriptions for *L. solani*, *L. equestris*, and *L. balteata*, respectively. Shaeffer (1919) suggested the possibility of synonymising *L. equestris* with *L. balteata* based on the morphological variability observed in a small percentage of *L. balteata* specimens, resulting in some phenotypes exhibiting a red prothorax and subapical spots. However, this synonymization was never completed due to a lack of rigorous comparison between species, and *L. balteata* remains a distinct species. When comparing specimens and images of the three species, the dominant phenotype of each species remains distinct within its species’ range. Hawaiian specimens typically exhibit the *L. equestris* phenotype (Figure 3) and, in combination with the molecular data, confirm the species’ identity. Rare intermediate specimens occur in Hawai‘i, exhibiting some characteristics of the *L. balteata* phenotype (Supplemental Figure 7C-D), but these phenotypes are unstable and should not be considered support for synonymising the species. While no specimens have been observed that exhibit all of the morphological characteristics expected of a species from another range (see Table 1 for these characteristics), observing the pattern of setae on the clypeus typically permits discrimination in intermediate specimens.

**Figure 2.**
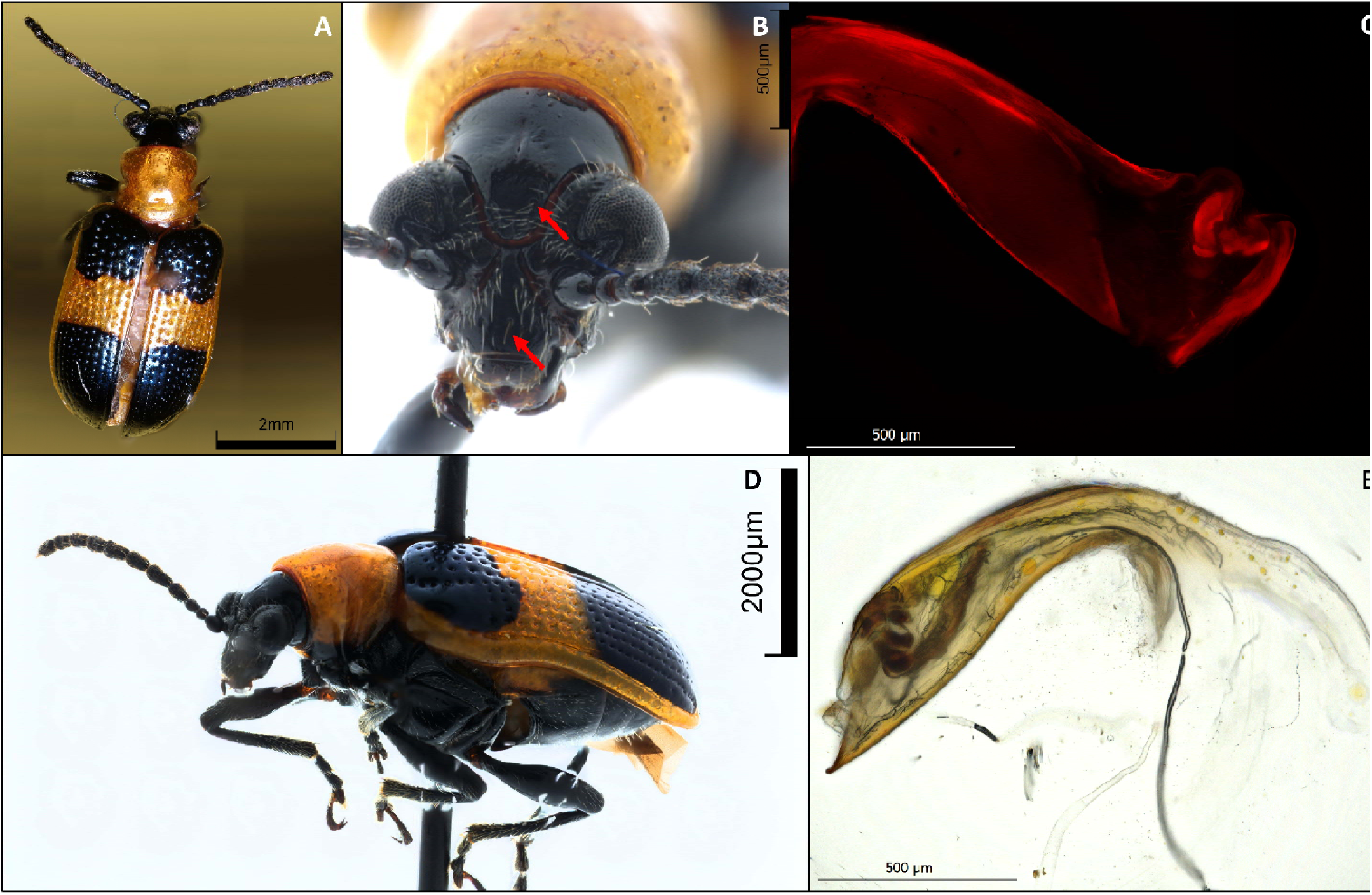
Images of *Lema solani*. A. Full body dorsal view. B. Dorsal view of the head, with red arrows showing diagnostic clypeal setae and setal pits on the frons that can be used to distinguish *L. solani* from *L. equestris*. C. Autofluorescent image of sclerotized structures of the aedeagus. D. Full body lateral view with a red arrow indicating the distinguishing unicolour elytral margin. E. Colour image of the aedeagus.

**Figure 3.**
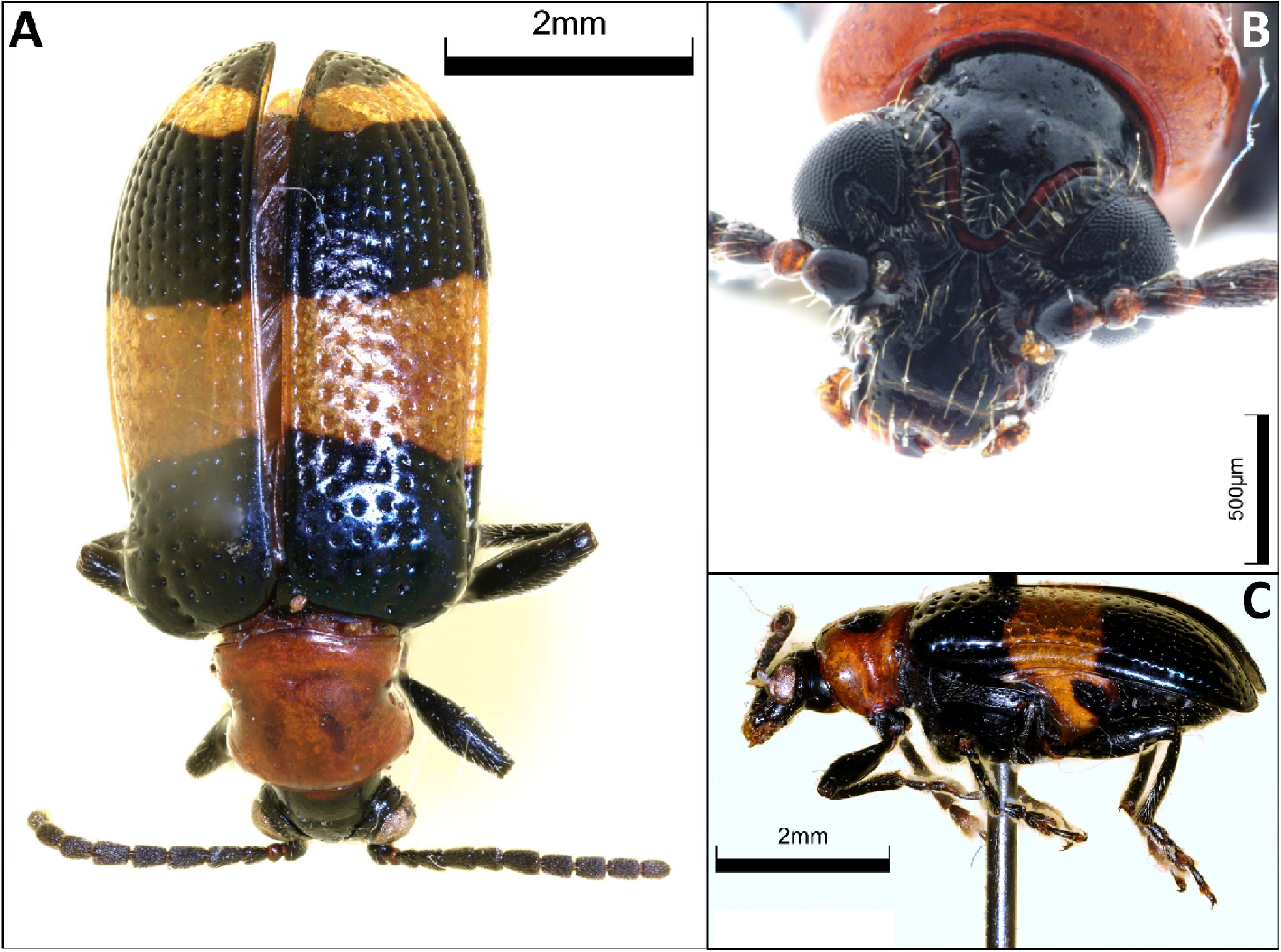
Images of *Lema equestris*. A. Full body dorsal image. B. A close-up of the head. C. Full body lateral image.

**Figure 4.**
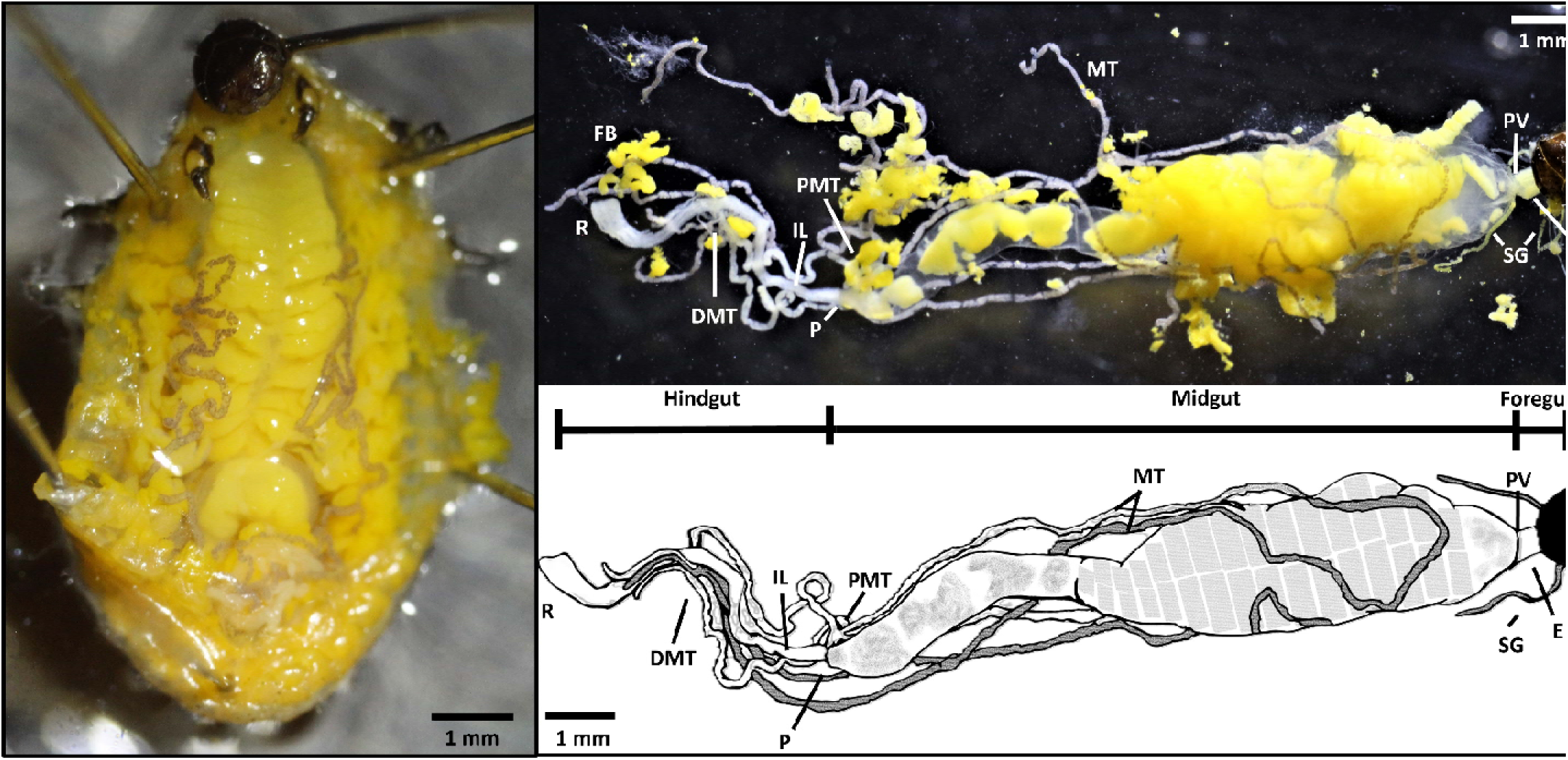
Photographs (left, top) and drawing (bottom) of the digestive system of *L. equestris* from an L4 larva. DMT. Distal malpighian tubule attachment point. E. Esophagus. FB. Fat Body. H. Head. IL. Ileum. MT. Malpighian tubules. P. Pylorus. PMT. Proximal malpighian tubule attachment point. PV. Proventriculus. R. Rectum. SG. Salivary glands.

**Table 1.**
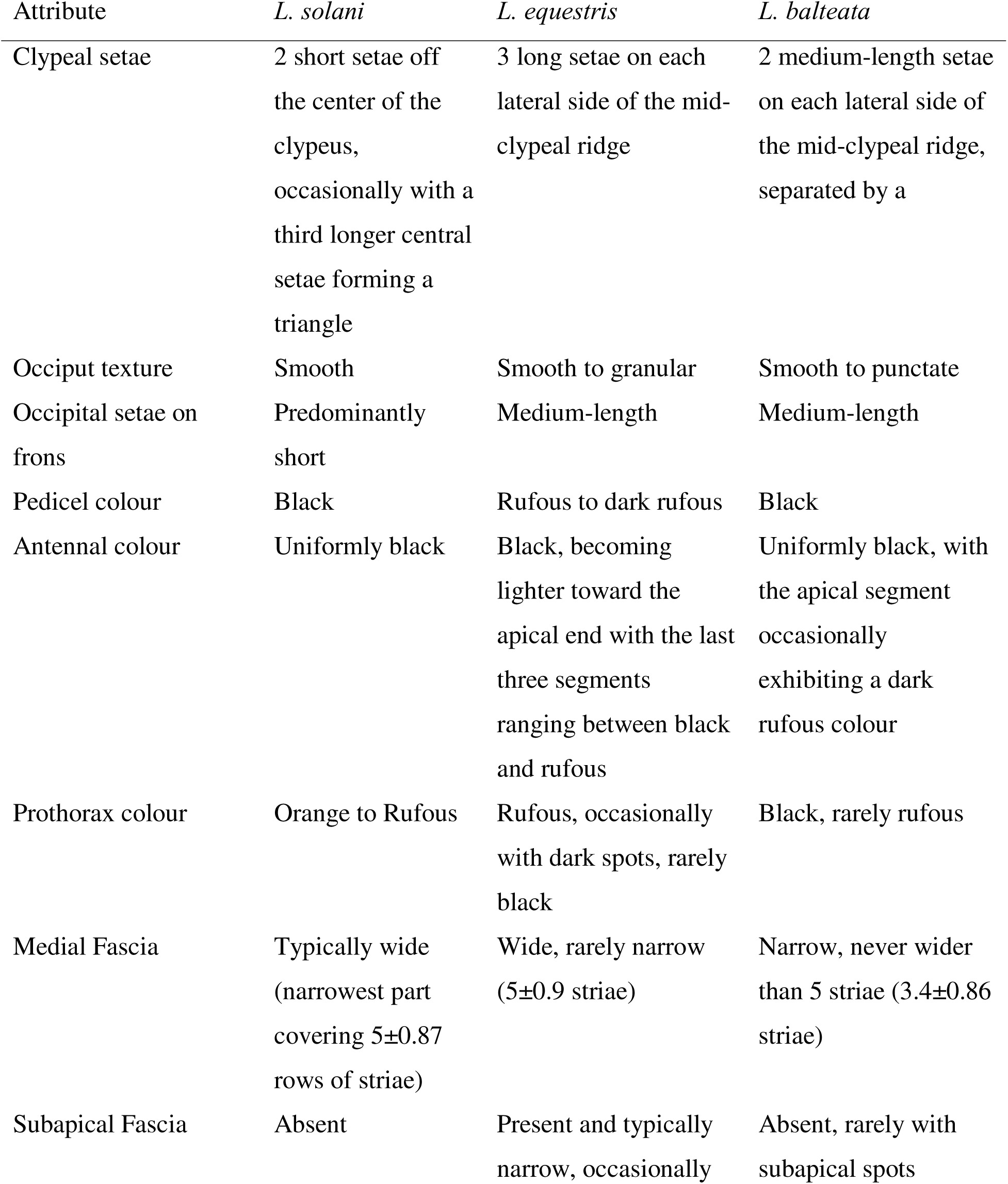
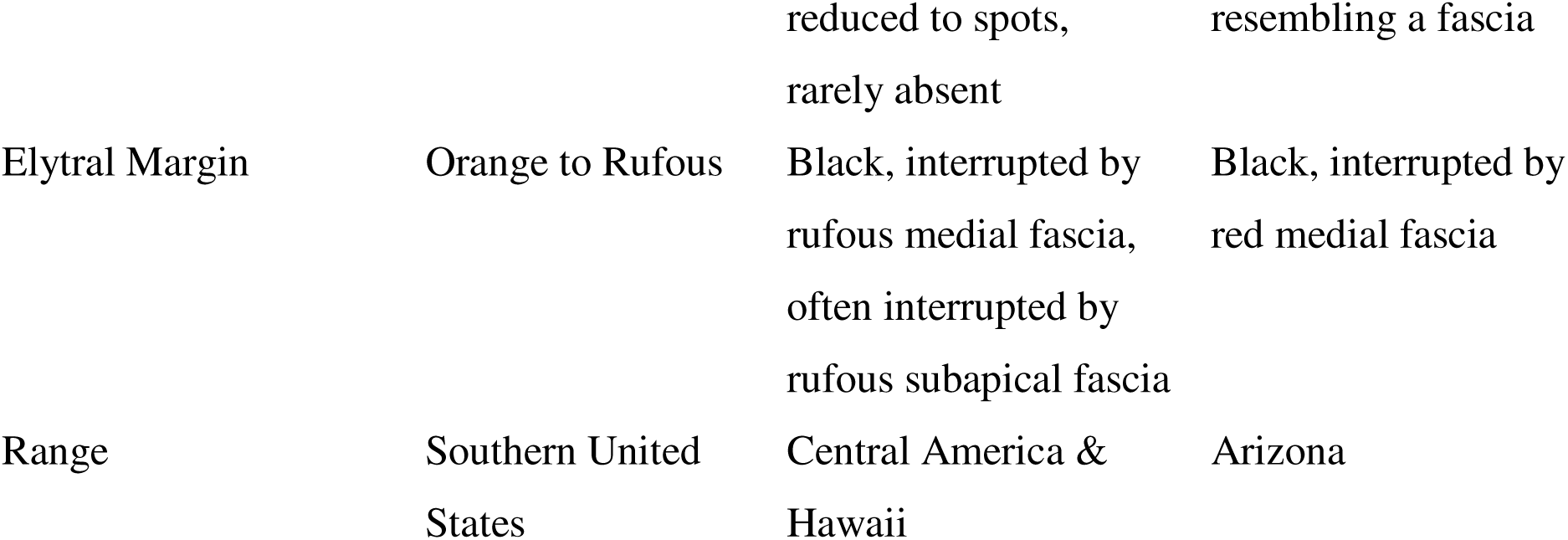
Discriminatory characteristics between *L. solani*, *L. equestris*, and *L. balteata*. Fascia width is reported as mean±SD, based on n=30 for each species.

The digestive anatomy of *L. equestris* aligns with other observations from Chrysomelidae (Özyurt Koçakoğlu *et al*., 2022). It has a shortened foregut, primarily composed of a muscular esophagus that ends in a small, conical proventriculus. After exiting the proventriculus, the food is enveloped by the peritrophic matrix, forming regularly shaped packets. The midgut accounts for the majority of the digestive anatomy, with large amounts of food being stored and digested in the anterior midgut.

The beetles have four malpighian tubules that branch off from a common trunk at the pylorus and end in a cryptonephridial complex where the tubules connect back to the hindgut at the distal attachment point in a common trunk and then run the length of the rectum (Figure 4). Two of the tubules are elongated and surround the anterior midgut, with a transition from white to yellowish brown as they reach the gut. The other two malpighian tubules are shortened and connect directly to the distal attachment point, staying white throughout their entire length. Larval and adult guts are similarly structured, with the primary differences being that the midgut is shortened by the absence of the anterior midgut storage and that all the malpighian tubules are white.

The reproductive anatomy closely resembles that of other *Lema* species found in the Americas, with similarly shaped spermathecae (Figure 5), aedeagi (Figure 6), and internal sclerites of the aedeagus (White, 1993; Monti *et al*., 2020). The spermatheca is a small, strongly sclerotized organ comprised of two chambers connected by a thinner tube and folded into a compact circular shape. The spermatheca connects, via the spermathecal duct, near the base of the large ovaries and oviduct. Males have two testes, which produce the filamentous sperm (Supplemental Figure 11), each composed of two testicular follicles and similar in size to the aedeagus. The internal sac of the aedeagus lacks a flagellum, agreeing with other *Lema* species within the *Quasilema* subgenus (Matsumura & Yoshizawa, 2012; Monti *et al*., 2020). Although similar, the structure of the internal sclerites of the aedeagus helps distinguish *L. equestris* from *L. solani*. In a ventral view focusing on the opening of the aedeagus, the ventral sclerite of *L. equestris* occludes the dorsal sclerite, hiding the two projections that visibly extend beyond the ventral sclerite in *L. solani*.

**Figure 5.**
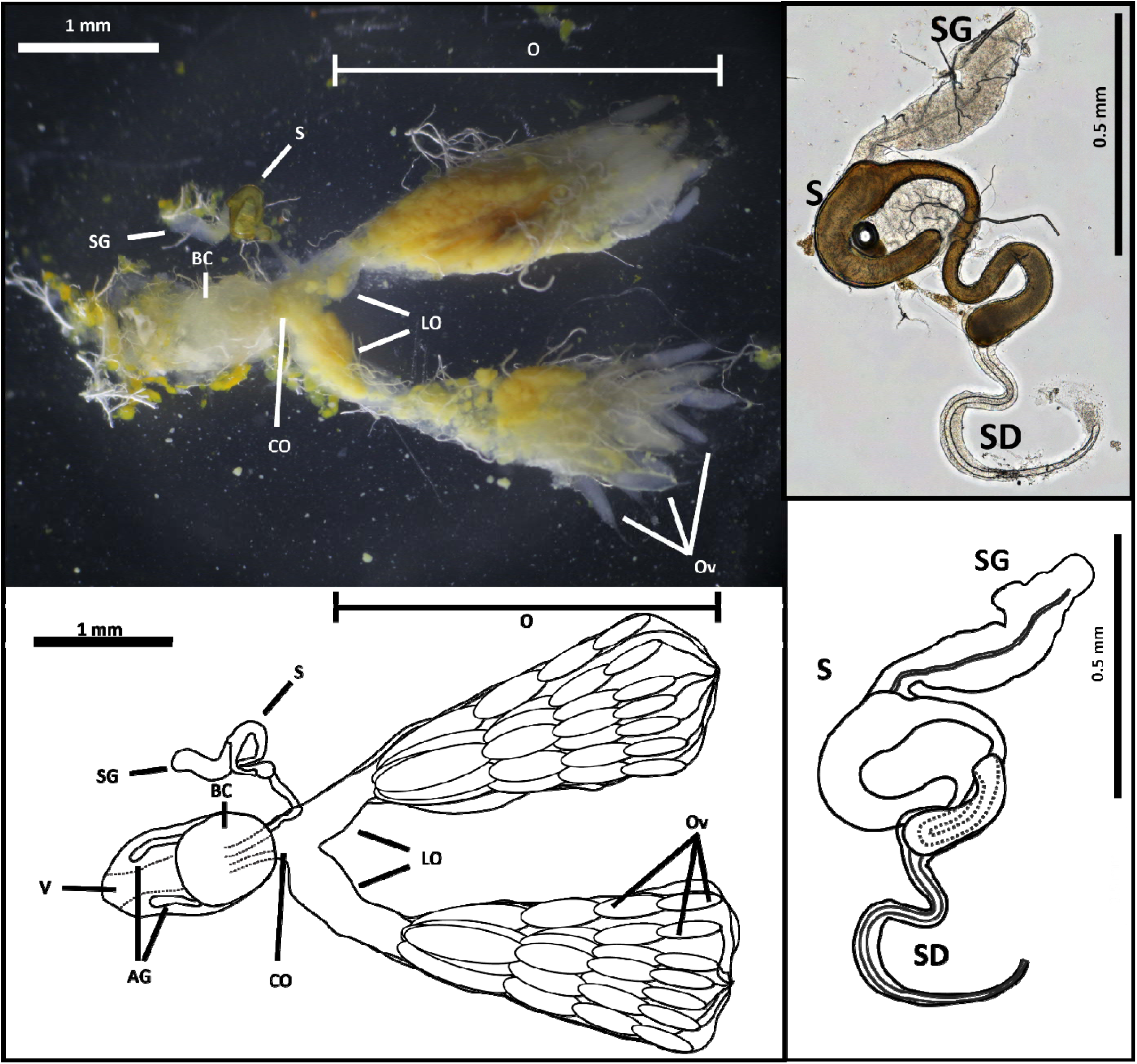
Photographs and drawings of the adult female reproductive system overall (left) and the spermatheca (right). The spermathecal micrograph is of a partially flattened specimen, while the drawing and the overall photograph show the specimen in its’ unbroken form.AG. Accessory glands (obscured in the photograph). CO. Common oviduct. LO. Lateral oviducts. O. Ovary. Ov. Ovarioles. S. Spermatheca. SD. Spermathecal duct. SG. Spermathecal gland. V. Vagina.

**Figure 6.**
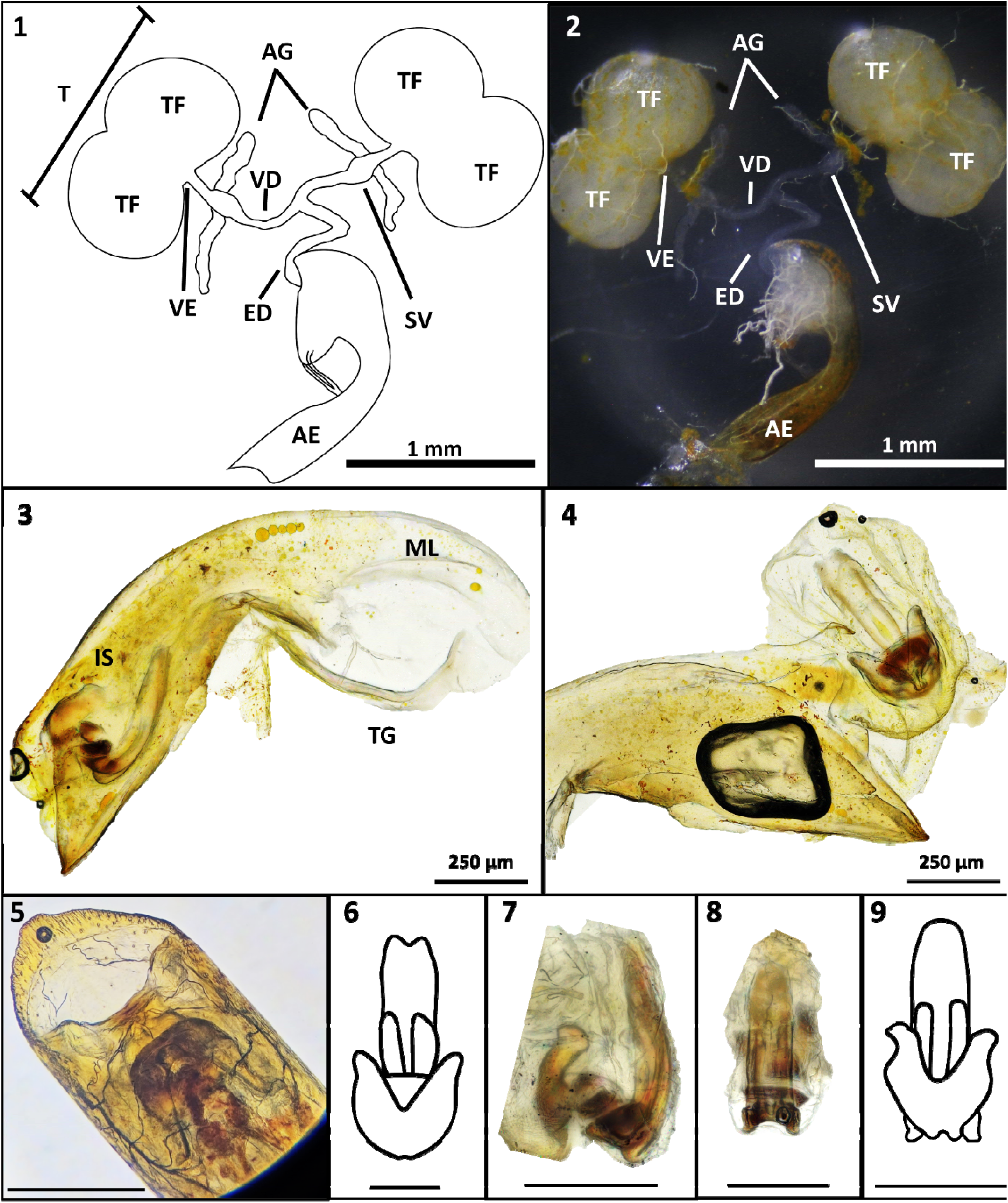
Drawing and photograph of the male reproductive system overall (1-2) with micrographs and drawings of the aedeagus and internal sclerites of *L. equestris* (3-6) and *L. solani* (7-9). Views are lateral (3, 6-7), dorsal (4, 5, 9), and ventral (8). AE. Aedeagus. AG. Paired accessory glands. ED. Ejaculatory duct. IS. Internal sclerites. ML. Median lobe. SV. Seminal vesicle. VD. Vas deferens. VE. Vas efferens. T. Testi. TF. Testicular follicle. TG. Tergum. Scale bars are equal to 250 microns unless stated otherwise.

### Rearing and Life Stages

Eggs of *L. equestris* are deposited along a rib, typically the midrib, on the underside of the leaves. Rearing efforts showed that female beetles would not oviposit on tissue paper, unlike other closely related beetles (Gelman *et al*., 2001). A count of 20 clutches revealed an average of 27 eggs per clutch with counts ranging from 13 to 48 (SD ±9.50). Freshly deposited eggs are pale yellow, with dark olive-coloured caps and a sticky, dark olive fluid that holds the base of the eggs to the leaf (Supplemental Figure 8). As the eggs mature, they darken from pale yellow to olive-yellow. Within 24 hours of hatching, typically around 7 days after the eggs are deposited, eye spots, movement, malpighian tubules, and two dark abdominal spots are visible through the chorion. Twelve hours prior to hatching, the tracheal system is visible, and the larvae contract, leaving an air space within the egg. Finally, within six hours of hatching, the larval head capsule will sclerotize, and larvae may begin moving within the egg in preparation for chewing through the chorion.

After hatching, *L. equestris* undergoes four larval instars. The middle two instars are short-lived, lasting only approximately one day in natural conditions with abundant food available (with a temperature and humidity of 25°C and 70%). The first and fourth instars are more variable, with one lasting one day and the other two days. Whether the first or fourth instar lasted two days varied by clutch, with all larvae in the same clutch following the same schedule. Sixty percent of larvae observed had a shortened first instar and a lengthened fourth instar (n=66). Identification of larval instar can be made using the following characteristics:

### L1

L1 larvae can be identified by their small size, reaching average length of 1.46±0.17 mm (n=43) just prior to molting, a lack of pronounced fat bodies visible through the cuticle, and two dark spots visible on the dorsolateral portion of the first abdominal segment. During this instar, larvae grow at a rate of 0.04 mm/hr.

### L2

L2 larvae are identifiable by their small size (2.42±0.27 mm, n = 22) and the beginnings of lateral fat body growth. Some fat may be present along the dorsal midpoint, but all the fat is still thin and appears dark yellow in colour due to the dark-coloured midgut being visible through the fat.

### L3

L3 larvae have developed thick layers of lateral fat that are orange to yellow in colour and no longer translucent. However, the fat bodies do not yet cover the entire dorsal surface, and the midgut is still visible through the cuticle.

### L4

L4 larvae are yellow in colour and are quite large (8.6±1.3 mm, min-max = 7-12 mm, n=26). The fat bodies on L4 larvae cover most to all the dorsal surface, obscuring the view of the midgut.

Final instar larvae move toward the ground by crawling downward along the plant’s stem or, if an obstacle is reached, dropping off the plant. Once on the ground, they exhibit negative phototaxis and move into debris or loose soil, where the larvae enter a prepupal stage. This prepupal stage is typically sessile but can move in response to light exposure over a moderate period (5-10 minutes). During the prepupal stage, the salivary glands undergo a transformation where they enlarge and become translucent (Supplemental Figure 9). This transition prepares the salivary glands to produce the foaming compound, which is then used to build the pupal cell (Supplemental Figure 10). The extent of pupal cell formation appears highly dependent on the larvae, with only a small amount of foam produced by larvae that transition to pupae early. This variance suggests that the amount of foam produced is directly linked to pupal energy reserves. While larvae generally look wrinkled at the prepupal stage, collection location and a lack of feeding are the only reliable external indicators. This prepupal stage can last up to 24 hours.

Pupae are orange in colour with translucent wings and antennae during early pupation. In the late pupal stage, a dark band will develop across the wings, and the eyes will darken. Pupae can wiggle when disturbed but otherwise remain in their pupal case until eclosion.

Adult beetles have a black head and black elytra with a metallic lustre visible at steep angles. Typically, the pronotum is red, and the black elytra have a medial and apical transverse fascia that intersects the elytral margin (Figure 3) and a stridulatory rake on the underside of the apical tip (Supplemental Figure 12). The elytral fascia can vary from rufous to orange (Supplemental Figure 7A-B), with rufous being the most common phenotype observed. While rare, the pronotum can have varying amounts of black, even to the point of appearing completely black dorsally (Supplemental Figure 7C-D). Specimens that have a darkened pronotum typically have a reduced apical elytral fascia and a larger portion of black on their abdomen.

Despite being capable fliers, adults may disperse more effectively when external factors facilitate it. Adult beetles typically fly short distances (<10 m) when disturbed or released and remain within the area. They can probably fly longer distances, but this has not been observed. It is believed that some percentage of adults typically remain within their natal grounds when food is still available. Supporting this, once a beetle population has been found, it is rarely extirpated if its host remains. Additionally, in areas with phenotypic abnormalities, such as black collars, beetles with similar abnormalities have been found months later at the same spot, but not in areas 100 m or more away.

Adults will continue to feed throughout their lives and mate repeatedly. In single-mate pairings, beetles have been observed repeatedly mating for up to 3 days when no opportunity for egg-laying has been provided. If an opportunity to lay eggs is provided, males will remain mounted for up to 24 hours after the initial mating before separating to lay eggs and then remating. When multiple males are nearby, other males have been observed remating the female within minutes of the first male dismounting.

### Diet Preference and Host Plant Trials

#### I. Adult feeding assays

Across all adult tests, adults regularly fed on *S. americanum* and almost no other plant (Figure 7). In the choice test, adults fed only on *S. americanum*, with no feeding or feeding damage observed on any other plants. Similarly, no feeding was observed on Brazilian nightshade, potato, tomato, eggplant, tomatillo, cabbage, cultivated tobacco, or tree tobacco and only minor damage was seen on poha berry. Poha feeding gave an expected rate of 0.01 mm^2^ beetle^−1^ h^−1^ (95% CI: [0.001, 0.042]), which was below the threshold for negligible adult feeding (P_p*mu < ROPE_ = 1.00) and 337-fold [64, 5139] less than on *S. americanum* (Expected rate: 3.32 mm^2^ beetle^−1^ h^−1^ [1.31, 5.45], P_p*mu < ROPE_ = 0.0006). Because adult beetles did not feed on any of the other plants, the computed probability of feeding was nearly identical (0.03 [0.00, 0.38]), with beetles more likely to feed on *S. americanum* (Difference_P_ = 0.74 [0.21, 0.98]). In the tests, beetles always fed on *S. americanum* when replacing test leaves, demonstrating that the absence of feeding resulted from host preferences rather than preexisting satiation.

**Figure 7.**
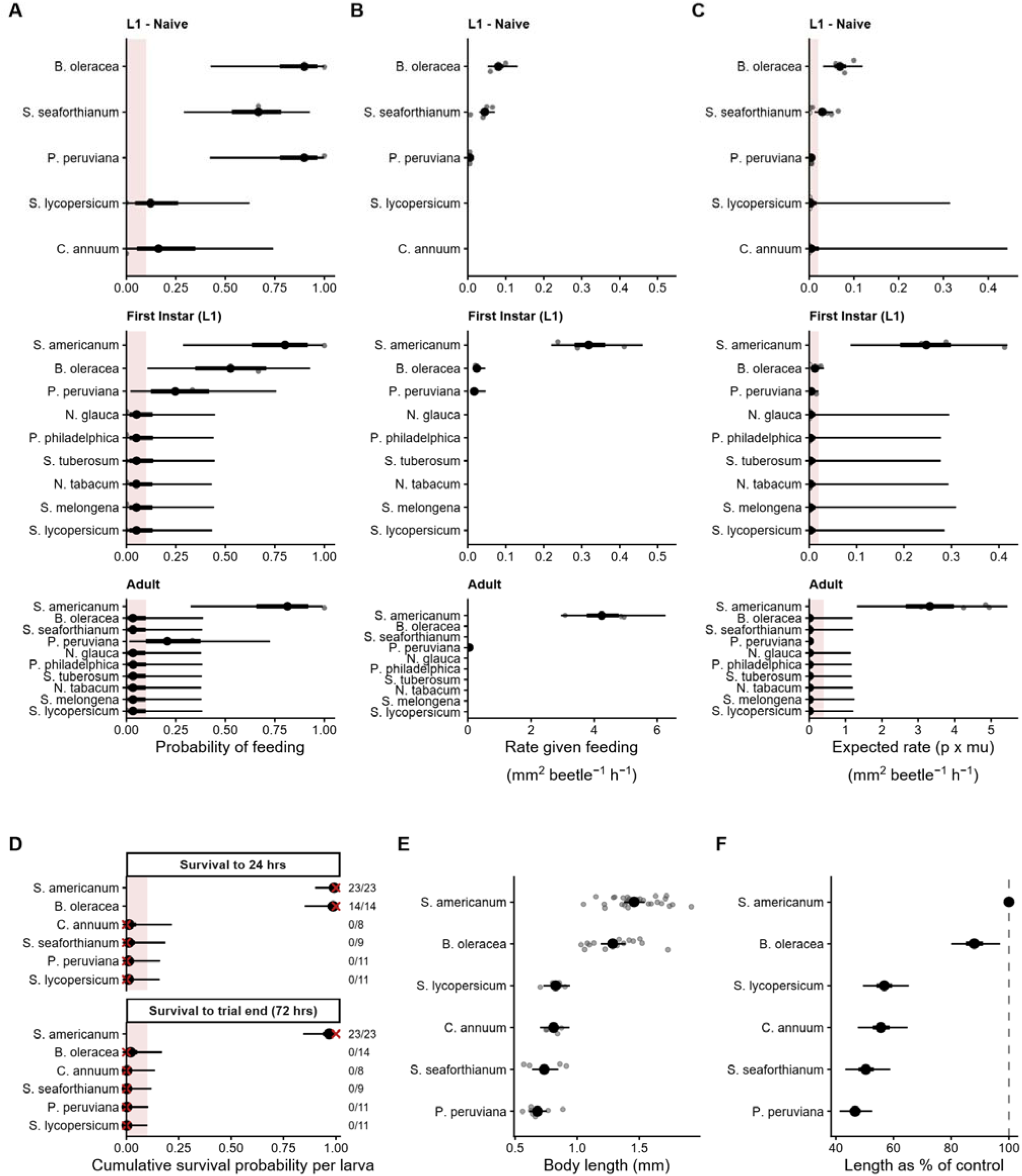
Posterior estimates for feeding (A-C), survival (D), and growth (E-F) of *L. equestris* beetles across tested plant species. Points represent the posterior median, with the bold and thin lines representing the 50 and 95% equal-tailed credible intervals, respectively. The red zone is the region of practical equivalence, wherein probabilities within this zone are biologically equivalent to zero. Raw data are shown as grey points. For survival probability, red “x” symbols and numbers denote the ratio of survivors to total larvae tested. Feeding estimates are separated by the probability that the beetles fed or not (A), the feeding rate provided that the beetles fed (B), and the feeding rate of beetles across all trials for the species, regardless of feeding status (C). Survival estimates (D) are separated into whether beetles would survive 24 hours or the full 72-hour trial. Posterior estimates of the body length are presented as length 24 hours after hatching (E), and the final length is calculated as a percentage of *S. americanum* (F).

#### II. Larval no-choice feeding assay

Larvae also showed a strong feeding preference for *S. americanum* (Figure 7). While larvae tasted the leaves of all species, most leaves had no detectable damage after four hours. Feeding was observed on *S. americanum* at an expected rate of 0.247 [0.087, 0.418] mm^2^ beetle^−1^ h^−1^, with prolonged biting on *B. oleracea* (2/3 trials) and *P. peruviana* (1/3 trials) lasting less than 30 minutes, after which the larvae roamed and sought a more suitable host. There was no feeding observed on any other species. While the probability that a beetle fed on cabbage was not distinguishable from *S. americanum* using the model (difference 0.23, 95% CI [-0.34, 0.76]; direction P = 0.80), the amount consumed once feeding began was 13.4-fold [6.2, 26.3] lower, and the expected rate 19.8-fold lower [5.5, 104.8]; difference 0.234 [0.074, 0.404]). Poha berry gave both a lower probability of any feeding (0.25 [0.02, 0.76]) and an expected rate 56-fold [9, 779] below S. americanum. While these feeding attempts demonstrated a strong food-seeking drive, none of the typical larvae tested in the 24-hour feeding trial grew, produced frass, or were found on the leaves at the end of the trial. This contrasted with the L1 larvae on the control plant, *S. americanum,* which doubled in size by the end of the 24 hours.

#### III. Larval host-naïve no-choice feeding assay

Host-naïve larvae were the most likely life stage to attempt feeding on plants besides *S. americanum* and showed a stronger feeding response to both cabbage and Brazilian nightshade than larvae that had already fed on *S. americanum*. Shortly after hatching, the naïve larvae attempted to feed on cabbage, poha berry, and Brazilian nightshade. Larvae fed at an expected rate of 0.069 [0.031, 0.119] mm^2^ beetle^−1^ h^−1^ on cabbage, Brazilian nightshade at 0.029 [0.012, 0.054], and negligibly on poha berry at 0.004 [0.002, 0.009]. The feeding rate on cabbage exceeded Brazilian nightshade by 2.36-fold [0.9, 6.5], and both exceeded poha berry (cabbage versus poha berry, ratio 15.4 [5.6, 39.7]; Brazilian nightshade versus poha berry, 6.5 [2.2, 17.2]), though all were dramatically less than control larvae (cabbage 3.5-fold less [1.17, 9.0]; Brazilian nightshade 8.32-fold less [2.55, 23.1]; and poha 54-fold less [17, 140]). Host-naïve larvae did not attempt to feed on chili pepper, or tomato.

Despite these feeding responses, plants other than *S. americanum* failed to support larval growth and survival. After hatching, cabbage was the only non-host plant on which host-naïve larvae fed enough to produce frass; however, the larvae exhibited the same wandering behaviour observed in previous tests and ultimately experienced stunted growth (88.1 % of control larval size regardless of cohort [80, 97]), resulting in a failure to molt into L2 larvae and no larvae surviving the entire 3 days (compared to zero mortality among the control larvae). Similarly, on the other plants tested, the naïve larvae quickly failed to thrive, and none lived beyond the 24-hour mark. The model estimated survival to the end of the trial at 0.968 [0.84,4 0.998] on the control and at 0.020 (95% CI [0.000, 0.170]) on cabbage and 0.003 to 0.004 (widest interval [0.000, 0.129]) on the remaining test plants. By the end of the test, larvae hatched on all other plants were 52% [45.4, 60.6] smaller than control larvae. Larvae that hatched on tomato and chili pepper continued to exhibit no feeding response and died within 24 hours of eclosing.

## DISCUSSION

The genus *Lema* is an extremely species-diverse group, with approximately 1,300 species described (Warchałowski, 2011), resulting in hosts spanning multiple plant families including, but not limited to, Asteraceae, Brassicaceae, Liliaceae, and Solanaceae (Clark, 2004). Some subgenera show increased host specificity. For example, the subgenus *Quasilema,* to which the *Lema* species studied here belong, is known to specialise on Solanaceae (Jolivet & Hawkeswood, 1995) with most species displaying a preference for their typical host’s genus, but willingly feeding on alternate plants from other genera within the same family when their ideal host is unavailable (Kogan & Goeden, 1970; Peschken & Johnson, 1979; Jolivet & Hawkeswood, 1995). Two species within *Quasilema* that resolved as closely related to *L. equestris* and *L. solani* in the CO1 phylogram*, L. daturaphila* Kogan & Goeden, 1970, and *L. bilineata* Germar, 1824, have become major agricultural pests due to their ability to feed from a wide range of genera of hosts beyond their preferred hosts (Cooper & Miles, 1951; Force, 1966; Kogan & Goeden, 1970).

Because these polyphagous beetles are closely related, it has been a concern that *L. equestris* may harm important plants in its new range or spread elsewhere, where it could cause more damage. In the diet trials, multiple solanaceous plants were tested from various genera and clades within *Solanum* (*S. seaforthianum, S. lycopersicum, S. melongena, P. philadelphica*, *P. peruviana, C. annuum, N. tabacum,* and *N. glauca*), and most showed no feeding response (47 total negative response trials across 16 plant-life stage combinations), with no alternative hosts supporting larval growth or survival. Rather, the results suggest that *L. equestris* exhibits a high degree of host specificity and may only feed on one *Solanum* minor clade at worst, and one *Solanum* species at best. Of the tested species, *S. americanum* and *S. seaforthianum* both belong to the DulMo major clade, with the former in the morelloid minor clade and the latter belonging to the dulcamaroid minor clade. Belonging to the potato major clade [potato and tomato minor clades, respectively] are *S. tuberosum* and *S. lycopersicum*, while *S. melongena* and *S. carolinense* are members of the leptostemonum major clade [eastern hemisphere spiny and carolinense minor clades, respectively].

Because larval *L. equestris* exclusively utilised *S. americanum,* a morelloid, it is likely that beetles do not pose a risk to endemic Hawaiian species of *Solanum,* which belong to different clades, or any other non-morelloid species. In Hawai‘i, members of the morelloid clade are occasionally grown and sold by native-plant nurseries and used as a traditional food or medicine. The importance of morelloids increases outside of Hawai‘i, however, with *S. americanum* grown as a vegetable crop in Indonesia (Mulyanto *et al*., 2018), *S. americanum* and *S. opacum* supporting native animals in Australia and New Zealand (Symon, 1979; Särkinen *et al*., 2018), and 12 endemic morelloids found across South America (Knapp & Särkinen, 2018; Knapp *et al*., 2020, 2023). Should *L. equestris* spread to any of these areas, it could pose a major threat to these important plants, proving more detrimental than its current status as a garden pest.

While this study provides important biological information, further work is needed. One limitation is that the study used only a small-scale caged assay, and the remaining laboratory assays used detached leaves, which can potentially alter feeding responses (ten Broeke *et al*., 2016; Dias *et al*., 2022) but remains a common test method for no-choice studies of Chrysomelids (Lake *et al*., 2015; Lommen *et al*., 2017). Open-field choice assays are often regarded as the ideal assay method for the most natural response (Briese, 2005; Schaffner *et al*., 2018) but the lack of caged-assay adult feeding responses beyond the typical host, or of equal larval feeding, growth, or survival on other hosts during no-choice tests strongly suggests the results would not differ. Another limitation is that only one species within the morelloid minor clade could be tested. Other morelloids should be tested to confirm whether host specificity is restricted to *S. americanum*, or if it extends to other morelloids, thereby providing more precise information about invasive potential and risk for other regions.

With the combination of all available data, and despite some morphological variability among Hawaiian specimens, *L. equestris* stands as the best possible identification of this new invasive in Hawai‘i. I have shown that *L. equestris* has strong host specificity, with adults, larvae, and host-naïve larvae either not feeding or not surviving on non-host plants. This work has attempted to gather basic biological and morphological data to support identification and further studies or management of this species in Hawai‘i and other ranges, should it continue to spread.

## Supporting information

Supplemental Methods and Images

## ACKNOWLEDGMENTS

I would like to thank Charles Mason at the US Department of Agriculture Agricultural Research Service Pacific Basin Agricultural Research Center for graciously providing access to their imaging microscope, Angela and David Holbrook for providing transportation and accommodation for some of the host collections, and Gary Matsuo for providing further host material. The author declares that no funds, grants, or other support were received during the preparation of this manuscript and that the author has no relevant financial or non-financial interests to disclose.

## REFERENCES

Abbott IA & Shimazu C (1985). The geographic origin of the plants most commonly used for medicine by Hawaiians. Journal of Ethnopharmacology 14, 213–222.

Arnott C, Atwood J, Bartlett R et al. (2021). Biosecurity in a Global Invasion Hotspot. In: Invasive Alien Species pp. 270–284. John Wiley & Sons, Ltd.

Austin KA & Rubinoff D (2024). Patterns of extinction across Hawaiian Lepidoptera offer lessons from a diverse, neglected, and vulnerable endemic fauna. Biodiversity and Conservation.

Baret S, Baider C, Kueffer C, Foxcroft LC & Lagabrielle E (2013). Threats to Paradise? Plant Invasions in Protected Areas of the Western Indian Ocean Islands. In: Plant Invasions in Protected Areas: Patterns, Problems and Challenges (eds LC Foxcroft, P Pyšek, DM Richardson & P Genovesi) pp. 423–447. Springer Netherlands, Dordrecht.

Briese DT (2005). Translating host-specificity test results into the real world: The need to harmonize the yin and yang of current testing procedures. Biological Control 35, 208–214.

Broeke CJM ten, Dicke M & Loon JJA van (2016). Feeding behavior and performance of Nasonovia ribisnigri on grafts, detached leaves, and leaf disks of resistant and susceptible lettuce. Entomologia Experimentalis et Applicata 159, 102–111.

Burbano E, Wright M, Bright DE & Vega FE (2011). New record for the coffee berry borer, Hypothenemus hampei, in Hawaii. Journal of Insect Science 11, 117.

Clark SM (2004). Host plants of leaf beetle species occurring in the United States and Canada: (Coleoptera: Megalopodidae, Orsodacnidae, Chrysomelidae, excluding Bruchinae). Coleopterists Society, Sacramento, CA.

Cooper O & Miles P (1951). On Lema Trilineata - a beetle closely resembling the tobacco slug, attacking the Cape gooseberry. South African Journal of Science 47, 330.

Costion CM, Kitalong AH & Holm T (2009). Plant endemism, rarity, and threat in Palau, Micronesia: a geographical checklist and preliminary Red List assessment. Micronesica 41, 131–164.

Cox PA & Elmqvist T (2000). Pollinator Extinction in the Pacific Islands. Conservation Biology 14, 1237–1239.

DeNitto GA, Cannon P, Eglitis A, Glaeser JA, Maffei H & Smith S (2015). Risk and pathway assessment for the introduction of exotic insects and pathogens that could affect Hawai’i’s native forests. General Technical Report, No. PSW-GTR-250. USDA Forest Service, Pacific Southwest Research Station.

Dias CR, Ataíde LMS, Meijer TT, Venzon M, Pallini A & Janssen A (2022). Phytophagous mite performance on intact plants and leaf discs with different defence levels. Entomologia Experimentalis et Applicata 170, 941–947.

Evenhuis NL & Miller SE (Eds) (2015). Records of the Hawaii Biological Survey for 2014 Part II: Index. Bishop Museum Occasional Paper. Bishop Museum Press, Honolulu, HI, U.S.A.

Feenstra KR & Clements DR (2008). Biology and Impacts of Pacific Island Invasive Species. 4. Verbesina encelioides, Golden Crownbeard (Magnoliopsida: Asteraceae). Pacific Science 62, 161–176.

Force DC (1966). Reactions of the Three-Lined Potato Beetle, Lema trilineata (Coleoptera: Chrysomelidae), to Its Host and Certain Nonhost Plants. Annals of the Entomological Society of America 59, 1112–1119.

Gelman DB, Bell RA, Liska LJ & Hu JS (2001). Artificial diets for rearing the Colorado potato beetle, Leptinotarsa decemlineata. Journal of Insect Science 1, 7.

Heenan PB, Tongatule HE & Panel W (2026). IUCN Red List Regional Assessment of the Flora of Niue Utilising Indigenous Local Knowledge and Botanical Information. Journal of the Royal Society of New Zealand 56, e70036.

Hoang DT, Chernomor O, Von Haeseler A, Minh BQ & Vinh LS (2018). UFBoot2: Improving the Ultrafast Bootstrap Approximation. Molecular Biology and Evolution 35, 518–522.

IUCN (2024). The IUCN Red List of Threatened Species. IUCN Red List of Threatened Species. Version 2024-2. Available from: https://www.iucnredlist.org/en [Accessed 3 January 2025].

Jolivet P & Hawkeswood TJ (1995). Host-plants of Chrysomelidae of the world: an essay about the relationships between the leaf-beetles and their food-plants. Backhuys, Leiden.

Jupiter S, Mangubhai S & Kingsford RT (2014). Conservation of Biodiversity in the Pacific Islands of Oceania: Challenges and Opportunities. Pacific Conservation Biology 20, 206–220.

Kalyaanamoorthy S, Minh BQ, Wong TKF, Von Haeseler A & Jermiin LS (2017). ModelFinder: fast model selection for accurate phylogenetic estimates. Nature Methods 14, 587–589.

Keppel G, Morrison C, Meyer J-Y & Boehmer HJ (2014). Isolated and vulnerable: the history and future of Pacific Island terrestrial biodiversity. Pacific Conservation Biology 20, 136–145.

Kleeck MJV & Holland BS (2018). Gut Check: Predatory Ecology of the Invasive Wrinkled Frog (Glandirana rugosa) in Hawai‘i. Pacific Science 72, 199–208.

Knapp S, Chiarini F, Cantero JJ & Barboza GE (2020). The Morelloid clade of Solanum L. (Solanaceae) in Argentina: nomenclatural changes, three new species and an updated key to all taxa. PhytoKeys 164, 33–66.

Knapp S & Särkinen T (2018). A new black nightshade (Morelloid clade, Solanum, Solanaceae) from the caatinga biome of north-eastern Brazil with a key to Brazilian morelloids. PhytoKeys 108, 1–12.

Knapp S, Särkinen T & Barboza GE (2023). A revision of the South American species of the Morelloid clade (Solanum L., Solanaceae). PhytoKeys 231, 1–342.

Kogan M & Goeden RD (1970). The Host-Plant Range of Lema trilineata daturaphila (Coleoptera: Chrysomelidae). Annals of the Entomological Society of America 63, 1175–1180.

Kumar L & Tehrany MS (2017). Climate change impacts on the threatened terrestrial vertebrates of the Pacific Islands. Scientific Reports 7, 5030.

Lacordaire T (1845). Monographie des coléoptères subpentamères de la famille des phytophages. C. Muquardt, Bruxelles.

Lake EC, Smith MC, Dray FA & Pratt PD (2015). Ecological host-range of *Lilioceris cheni* (Coleoptera: Chrysomelidae), a biological control agent of *Dioscorea bulbifera*. Biological Control 85, 18–24.

Lee C-F & Matsumura Y (2013). On newly and recently recorded species of the genus Lema Fabricius (Coleoptera, Chrysomelidae, Criocerinae) from Taiwan. ZooKeys 262, 17–37.

Lommen STE, Jolidon EF, Sun Y, Bustamante Eduardo JI & Müller-Schärer H (2017). An early suitability assessment of two exotic Ophraella species (Coleoptera: Chrysomelidae) for biological control of invasive ragweed in Europe. EJE 114, 160–169.

Mason C, Weaver M, Kissinger K et al. (2026). Applying PCR cycle autonormalization to improve PacBio full-length 16S rRNA sequencing.

Matsumura Y & Yoshizawa K (2012). Homology of the internal sac components in the leaf beetle subfamily criocerinae and evolutionary novelties related to the extremely elongated flagellum. Journal of Morphology 273, 507–518.

McClelland DHR, Nee M & Knapp S (2020). New names and status for Pacific spiny species of Solanum (Solanaceae, subgenus Leptostemonum Bitter; the Leptostemonum Clade). PhytoKeys 145, 1–36.

Meyer JY (2004). Threat of Invasive Alien Plants to Native Flora and Forest Vegetation of Eastern Polynesia. Pacific Science 58, 357–375.

Monti MM, Ruocco M, Grobbelaar E & Pedata PA (2020). Morphological and Molecular Characterization of Lema bilineata (Germar), a New Alien Invasive Leaf Beetle for Europe, with Notes on the Related Species Lema daturaphila Kogan & Goeden. Insects 11, 295.

Mulyanto D, Iskandar J, Abdoellah OS, Iskandar BS, Riawanti S & Partasasmita R (2018). Leunca (Solanum americanum Mill.): The uses as vegetable in two villages in Upper Citarum Area, Bandung, West Java, Indonesia. Biodiversitas Journal of Biological Diversity 19, 1941–1954.

Münster P & Wieczorek AM (2007). Potential gene flow from agricultural crops to native plant relatives in the Hawaiian Islands. Agriculture, Ecosystems & Environment 119, 1–10.

Özyurt Koçakoğlu N, Candan S & Güllü M (2022). Anatomy and histology of digestive tract in the red poplar leaf beetle Chrysomela populi Linnaeus, 1758 (Coleoptera: Chrysomelidae). International Journal of Tropical Insect Science 42, 927–939.

Paradis E (2012). Analysis of Phylogenetics and Evolution with R. Springer New York, New York, NY.

Paudel S, Jackson TA, Mansfield S, Ero M, Moore A & Marshall SDG (2023). Use of pheromones for monitoring and control strategies of coconut rhinoceros beetle (*Oryctes rhinoceros*): A review. Crop Protection 174, 106400.

Peck RW, Mikros D, Dunkle EJ, Jaenecke K & Roy K (2026). Temporal associations between ambrosia beetles and ōhi a (Metrosideros polymorpha) artificially inoculated with Ceratocystis lukuohia. Agricultural and Forest Entomology 28, 49–60.

Peschken DP & Johnson GR (1979). Host Specificity and Suitability of Lema cyanella (Coleoptera: Chrysomelidae), a Candidate for the Biological Control of Canada Thistle (Cirsium arvense). The Canadian Entomologist 111, 1059–1068.

Reichard SH & White P (2001). Horticulture as a Pathway of Invasive Plant Introductions in the United States: Most invasive plants have been introduced for horticultural use by nurseries, botanical gardens, and individuals. BioScience 51, 103–113.

Rønsted N, Walsh SK, Clark M et al. (2022). Extinction risk of the endemic vascular flora of Kauai, Hawaii, based on IUCN assessments. Conservation Biology 36, e13896.

Särkinen T, Poczai P, Barboza GE, Weerden GM van der, Baden M & Knapp S (2018). A revision of the Old World Black Nightshades (Morelloid clade of Solanum L., Solanaceae). PhytoKeys 1–223.

Schaffner U, Smith L & Cristofaro M (2018). A review of open-field host range testing to evaluate non-target use by herbivorous biological control candidates. BioControl 63, 405–416.

Schindelin J, Arganda-Carreras I, Frise E et al. (2012). Fiji: an open-source platform for biological-image analysis. Nature Methods 9, 676–682.

Shaeffer C (1919). Synonymical and Other Notes on Some Species of the Family Chrysomelidæ and Descriptions of New Species. Journal of the New York Entomological Society 27, 307–340.

Symon DE (1979). Fruit Diversity and Dispersal in Solanum in Australia. Journal of the Adelaide Botanic Garden 1, 321–331.

Taylor L & Kumar S (2016). Global Climate Change Impacts on Pacific Islands Terrestrial Biodiversity: A Review. Tropical Conservation Science.

Trifinopoulos J, Nguyen L-T, von Haeseler A & Minh BQ (2016). W-IQ-TREE: a fast online phylogenetic tool for maximum likelihood analysis. Nucleic Acids Research 44, W232–W235.

USFWS (2019). Initiation of 5-Year Status Reviews for 91 Species in Oregon, Washington, Hawaii, and American Samoa. Notice, No. 2019–12244.

Van Driesche RG & Murray TJ (2004). Chapter 7. Overview of Testing Schemes and Designs Used to Estimate Host Ranges. In: Assessing Host Ranges of Parasitoids and Predators Used for Classical Biological Control: A Guide To Best Practice (eds RG Van Driesche & R Reardon) pp. 68–89. USDA Forest Service - Forest Health Technology Enterprise Team, Morgantown, West Virginia.

Villalobos EM, Nikaido S, Ito T & Wong J (2024). Monitoring strategies during the establishment phase of Aethina tumida on Oahu, Hawaii. Journal of Applied Entomology 148, 708–711.

Waldren S, Florence J & Chepstow-Lusty AJ (1995). Rare and endemic vascular plants of the Pitcairn Islands, south-central Pacific Ocean: A conservation appraisal. Biological Conservation 74, 83–98.

Warchałowski A (2011). An introductory review of Lema Fabr. species from Eastern and Southeastern Asia (Coleoptera: Chrysomelidae: Criocerinae). Genus. International Journal of Invertebrate Taxonomy 22, 29–93.

White R (1993). A revision of the subfamily Criocerinae (Chrysomelidae) of North America North of Mexico. Technical Bulletin - United States Department of Agriculture 1805, 43–44.

Wood KR, Oppenheimer H & Keir M (2019). A checklist of endemic Hawaiian vascular plant taxa that are considered possibly extinct in the wild. Technical Report, No. 314. The National Tropical Botanical Garden.

