## Supplemental Methods and Images for "A Characterisation of the Invasive Lema Beetle, *Lema equestris* (Coleoptera: Chrysomelidae), in Hawai‘i"

Mikinley Weaver[1,2]

[1] *The Center for Understanding Biological Sciences, Hilo, HI, USA*
[2] *Victoria University of Wellington, Wellington, New Zealand*

### 1. Bayesian Statistical Analysis

The following sections describe the analysis in further detail.

#### 1.1 Data structure

Each feeding trial $_i$ recorded whether a replicate of $n_i$ beetles, sharing a life-stage-by-plant combination g(i), consumed any of the plant over a recorded duration $H_i$ (h). When the hurdle was overcome, or when consumption was recorded ($fed_i = 1$), the leaf area consumed, $Y_i$, was used as the response variable. Across 24 life-stage-plant combinations, 75 trials were used in the final model, of which 21 recorded feeding and 54 did not and 16 of the 24 combinations never fed.

#### 1.2 Model

Whether a dish fed at all was modelled as a Bernoulli outcome at the level of the dish:

$$fed_i \sim \text{Bernoulli}(p_{g(i)})$$

$$\text{logit}(p_g) = \text{logit\_p\_mu}[s(g)] + \text{logit\_p\_z}_g \times \text{logit\_p\_sigma}$$

logit_p_mu[s] ~ normal(mean = 0, standard deviation = 2); logit_p_sigma ~ half-normal(mean = 0, standard deviation = 1); $\text{logit\_p\_z}_g$ ~ normal(mean = 0, standard deviation = 1) **(S1)**

The probability that a dish of beetles in combination g shows any feeding damage in a trial of the kind conducted is $p_g$, and s(g) is the life stage of that combination. $\text{logit\_p}_g$ is decomposed into a stage mean, a between-combination standard deviation and a standardised per-combination deviation (a non-centred parameterisation) to keep sampling stable across all combinations. When a dish fed, the amount consumed was modelled as:

$$Y_i \sim \text{gamma}(\text{shape} = \text{phi} \times n_i, \text{rate} = \text{phi} / (\text{mu}_{g(i)} \times H_i))$$

$$\log(\text{mu}_g) = \text{log\_mu\_mu}[s(g)] + \text{log\_mu\_z}_g \times \text{log\_mu\_sigma}$$

log_mu_mu[s] ~ normal(mean = log(0.1), standard deviation = 2); log_mu_sigma ~ half-normal(mean = 0, standard deviation = 1); $\text{log\_mu\_z}_g$ ~ normal(mean = 0, standard deviation = 1); phi ~ exponential(rate = 1) **(S2)**

The mean of $Y_i$ is $\text{mu}_{g(i)} \times n_i \times H_i$, so $\text{mu}_g$ is the mean consumption per beetle per hour of exposure given that the dish fed, and the coefficient of variation is $1/\text{sqrt}(\text{phi} \times n_i)$. Scaling the shape by $n_i$ is the distribution of a sum of $n_i$ per-beetle contributions, which is the assumption in S1. Exposure enters the mean linearly in both the number of beetles and the duration.

$$rate_g = p_g \times mu_g \quad \textbf{(S3)}$$

The quantity reported alongside the two components is their product, the unconditional expected consumption per beetle per hour (S3), estimated by the raw per-beetle-per-hour means. Of these parameters $p_g$, $mu_g$ and $rate_g$ are of primary interest, while logit_p_mu[s], logit_p_sigma, log_mu_mu[s], log_mu_sigma and phi govern the hierarchical structure and are not compared. Log_mu_mu was centred on the order of magnitude of per-beetle-per-hour consumption visible in exploratory plots of the raw data before fitting, with a standard deviation of 2 on the log scale, which spans roughly two orders of magnitude either side of that centre. The two half-normal(0, 1) scale priors are deliberate regularisers rather than uninformative choices: 16 of the 24 combinations never fed, so their group-level probabilities are bounded above by the data but not below, and logit_p_sigma is only weakly identified from above.

Survival was modelled as a staged process, because a larva that survived to the end of the trial had already survived to 24 h. Thus, for life stage k, replicate r, and plant g(r), writing m[k,r] for the number of larvae at the start of that stage, so that m[1,r] = n_placed[r] and m[2,r] = successes[1,r].

successes[k,r] ~ binomial(m[k,r], inverse-logit(logit_prob[k,g(r)]))

logit_prob[k,g] = bin_mu[k] + bin_z[k,g] x bin_sigma[k]

bin_mu[k] ~ normal(0, 1.5); bin_sigma[k] ~ half-normal(0, 1.5); bin_z[k,g] ~ normal(0, 1) **(S4)**

The probability of surviving to a time point is the product of the stages. The model provides both the cumulative probability and the stage-conditional probability. Final body length was modelled on the log scale, so that a plant-versus-control contrast is a ratio:

length[j] ~ lognormal(log_len[g(j)], len_resid)

log_len[g] = len_mu + len_z[g] x len_sigma_plant; len_mu ~ normal(log(1.5), 0.5);

len_sigma_plant ~ half-normal(0, 0.5); len_resid ~ half-normal(0, 0.25) **(S5)**

The reported quantity is the modelled mean length on a plant as a percentage of the modelled mean length of the control larvae:

length_pct_of_control[g] = 100 x exp(log_len[g] - log_len[control]) **(S6)**

### 1.3 Prior predictive check

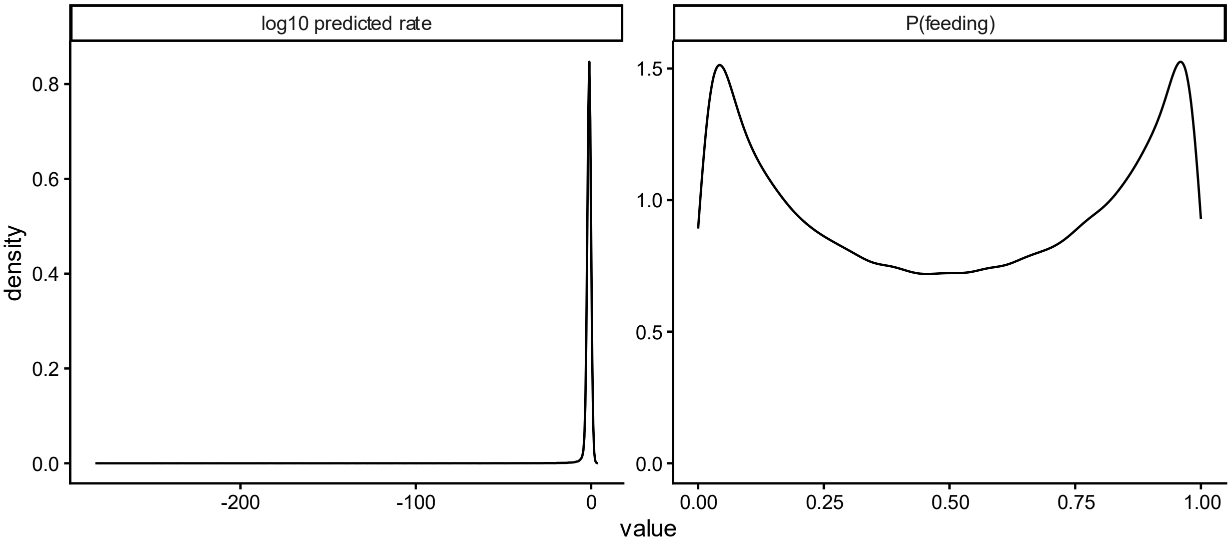


Supplemental Figure 1. Plotted parameter estimates for the feeding model with the predicted feeding rate on log scale (left) and probability of feeding (right).

Before fitting, the model was run with the likelihood switched off, so that the prior predictive check exercises the same code as the fit rather than a separate simulation that could drift from it. Simulated feeding probabilities spanned the full range from near-certain feeding to near-total refusal. The implied prior is consistent with the expectation that a plant is either a host or is not, and is not a flat prior on the probability scale (Supp. Fig. 1). Simulated positive consumption was strictly positive and right-skewed, spanning several orders of magnitude around the rates seen in comparable assays. For the development model checks, the survival probability prior spanned the entire probability range, but skewed toward non-survival while length was positive and left-skewed (Supp. Fig. 2).

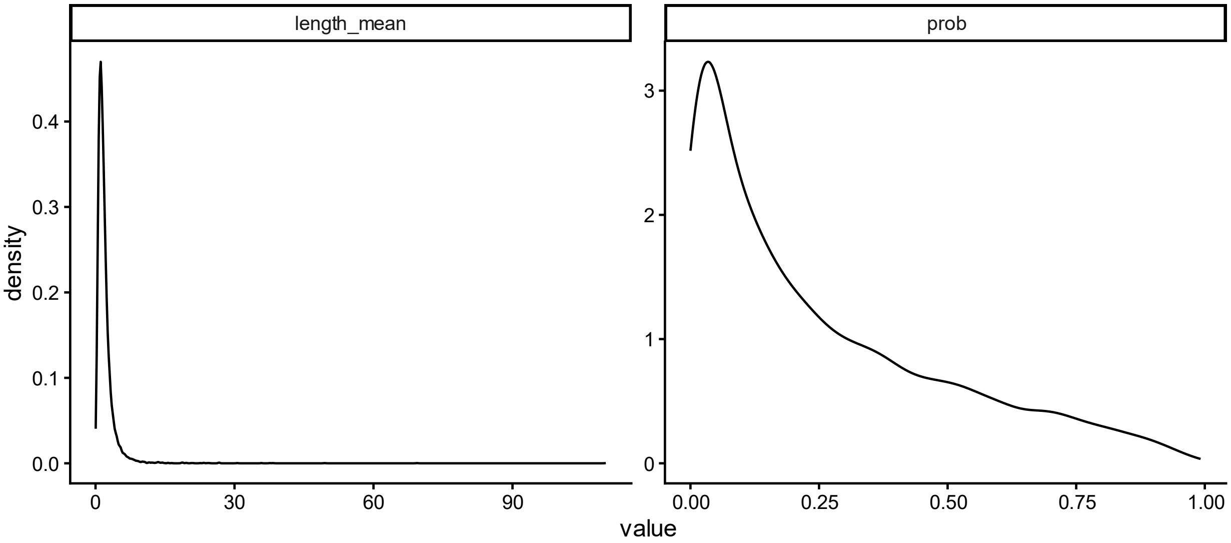

Supplemental Figure 2. Plotted parameter estimates for the development models with the predicted length (left) and the cumulative probability of survival (right).

### 1.4 Posterior predictive check

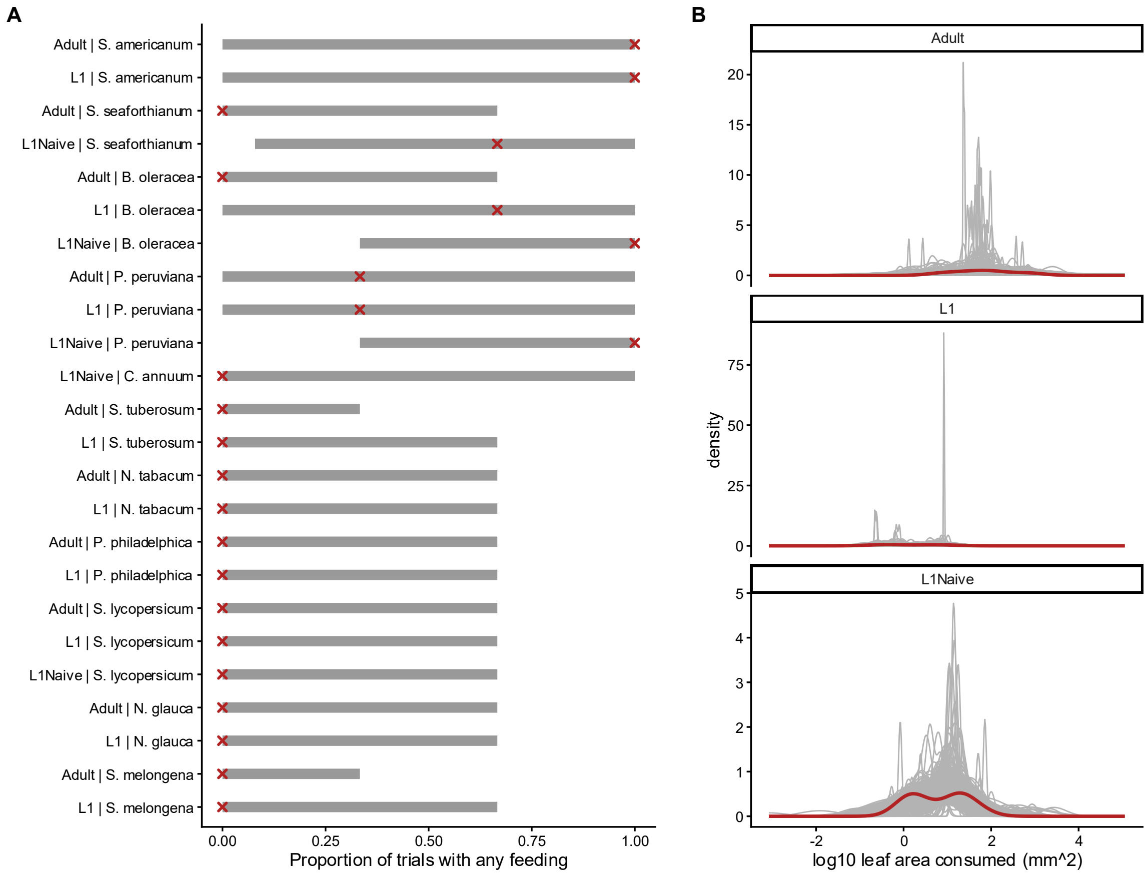


Supplemental Figure 3. Posterior predictive checks for the feeding model. A. Proportion of trials with any feeding, by life-stage-plant combination: observed values (red) against the simulated 95% intervals (grey). B. Distribution of positive consumption on the log scale, observed (red) and posterior-predictive replicates (grey).

The fitted model's ability to reproduce the data was assessed against replicate datasets generated inside the model itself, from 500 posterior draws, so that the check compares the fitted model with the data. For the hurdle component, the observed proportion of trials with any feeding was compared, for each life-stage-plant combination, against the simulated 95% interval of replicate proportions. For the amount component, the observed distribution of positive log-transformed consumption was compared against the posterior-predictive replicates, separately for each life stage as well as pooled. For both checks the observed values fell within expectations (Supplemental Figure S3).

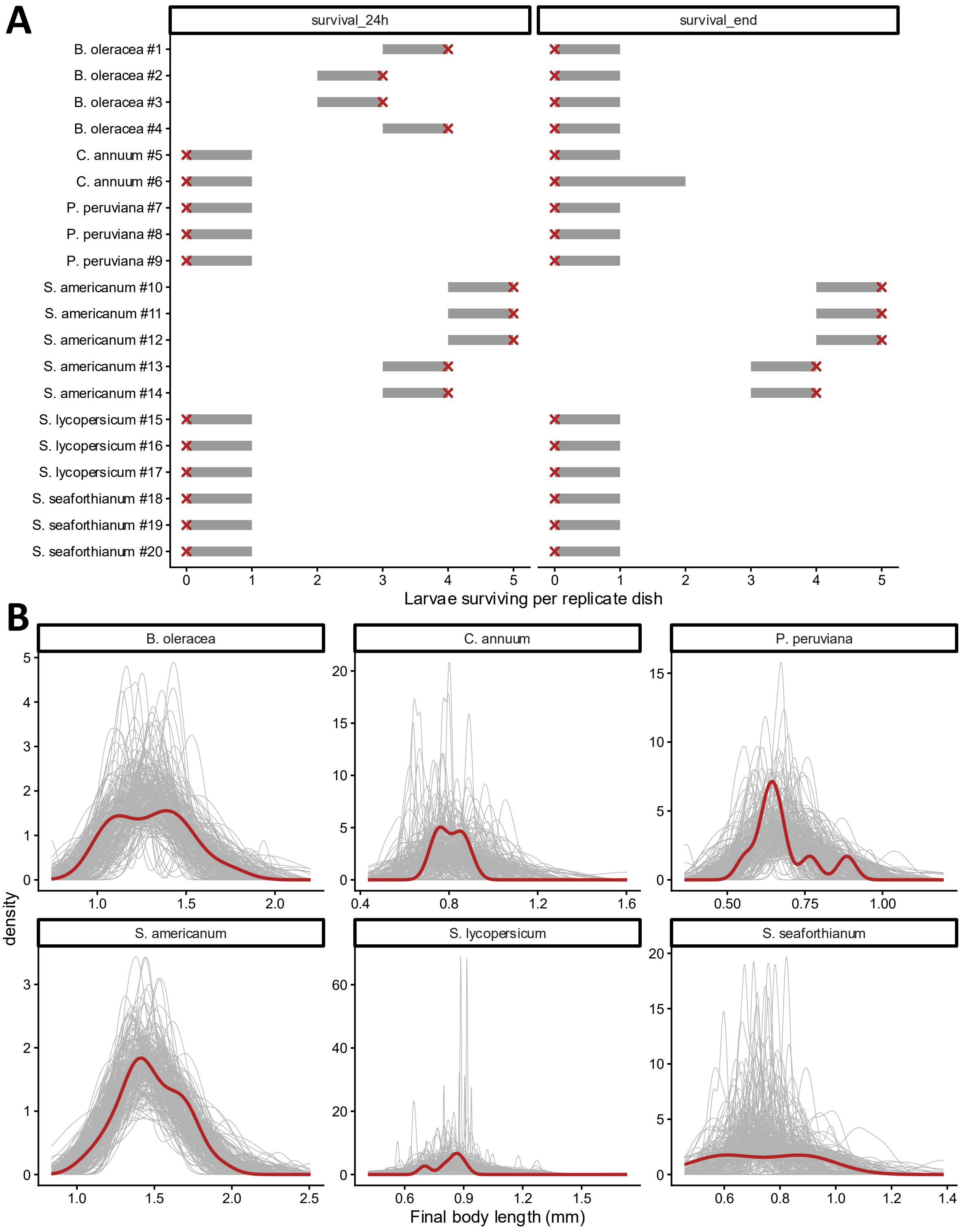


Supplemental Figure 4. Posterior predictive check for the survival and growth model. A. Observed survivors per replicate dish (red) against the simulated 95% intervals (grey). B. Observed distribution of final body length (red) overlaid on posterior-predictive replicates (grey), faceted by plant.

1.5 Contrasts

For every pair of plant species tested at the same beetle life stage, and for each of $p_g$, $mu_g$ and $rate_g$, both an absolute difference and a proportional ratio were calculated:

$$Difference_{g1,g2} = \theta_{g1} - \theta_{g2}$$

$$Proportion_{g1,g2} = \theta_{g1} / \theta_{g2} \quad \textbf{(S7)}$$

Contrasts of $mu_g$ were restricted to plant pairs in which feeding was observed at least once for both members. Contrasts of $p_g$ used every pair, since a probability is informed by zero counts. Ratios are formed as the value for the earlier plant of a pair divided by the value for the later one, and are reported only where both plants were fed on at least once. Contrasts of $rate_g$ are computed for every pair, but those in which one plant was never fed on are upper bounds, because one half of such a contrast carries a partially pooled $mu_g$ rather than an estimate of that plant.

1.6 Region of practical equivalence

The claim under test in these assays is a null one, that a plant is not a host. As described in the manuscript, regions of practical equivalence were specified before fitting.

$$P\_ROPE_{,g} = Pr(p_g \times mu_g < ROPE\_s(g)) \quad \textbf{(S8)}$$

Over a trial of the duration each life stage received, the three limits correspond to roughly 9.6, 0.12 $mm^2$ of leaf per beetle. For contrasts of $p_g$, which are on the probability scale and to which an area threshold does not apply, the region is a difference of less than 0.10: fewer 10% differing in outcome.

1.7 Sensitivity to the priors

Prior constants are supplied to the model as data, so each sensitivity condition refits the same model file. Each of the six prior constants was halved and doubled in turn, and the centre of the prior on log_mu_mu was moved an order of magnitude in each direction, giving thirteen fits including the reported one. Estimates of consumption were essentially unchanged across every condition, except those where feeding was unobserved, and no fit produced a divergent transition. Feeding probabilities for combinations in which feeding was never observed are the exception as they move with the prior on the between-combination standard deviation, and rather than as measurements. This behaviour is expected, should be read as regularised upper bounds, and is the reason the prior-posterior comparison (Supp. Fig. 5) shows the two scale parameters (logit_p_sigma, phi) lying in the upper tails of their priors.

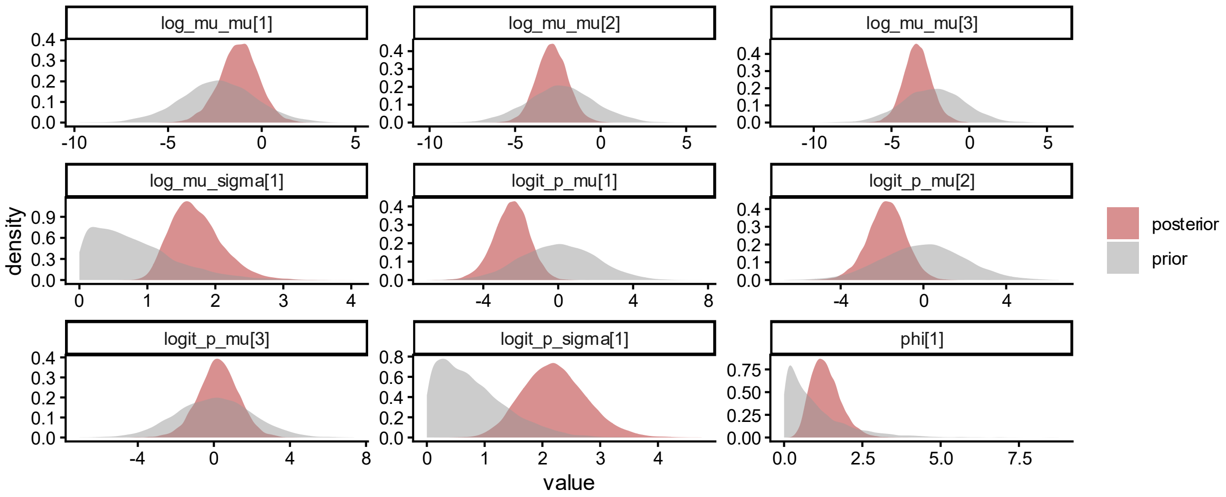


Supplemental Figure 5. Prior-posterior comparison across the feeding model parameters.

### 1.10 Influential observations

Approximate leave-one-out cross-validation, with Pareto-smoothed importance sampling, was used to check whether any single trial carries an estimate. Two trials with a Pareto k above 0.7 were identified, originating from the adult and L1 trials with poha. These combinations flagging here are a result of each consisting of beetles that fed in only one of three replicates.

### 1.11 Predicted feeding rate

To visualise feeding rate, posterior predictions were generated for each combination:

$$fed_{pred} \sim \text{bernoulli}(\text{probability} = p_g);\ \ rate_{pred} = fed_{pred} \times Y_{pred} / (n_g \times H_g),\ \ Y_{pred} \sim \text{gamma}(\text{shape} = phi \times n_g,\ \text{rate} = phi / (mu_g \times H_g)) \quad \textbf{(S9)}$$

The prediction combines both sources of uncertainty, whether feeding occurs at all and how much occurs given that it does, into a single predictive distribution per combination, expressed per beetle per hour so that the observed trial rates overlay it directly. This predictive distribution includes the Bernoulli outcome and the gamma observation noise, as well as uncertainty in the parameters, so it is wider than the posterior for the expected rate.

Supplemental Table S1. GenBank accession numbers of sequences used.

| | | |
|---|---|---|
| LEMA sp. | | |
| HQ582576.1 | HQ582645.1 | AY242400.1 |
| MT456399.1 | MT456400.1 | PV990476.1 |
| MT456401.1 | | |

### LEMA CONCINNIPENNIS

| | | |
|---|---|---|
| OL663182.1 | OL663183.1 | OL663184.1 |

### LEMA DECUMPUNCTATA

| | | |
|---|---|---|
| OL663191.1 | OL663192.1 | OL663193.1 |

### LEMA DATURAPHILA

| | | |
|---|---|---|
| HM431490.1 | HM431491.1 | MT002938.1 |
| HM431494.1 | KR481201.1 | MT002940.1 |
| MF640993.1 | MT002939.1 | |

### LEMA CYANELLA

| | | |
|---|---|---|
| JF889862.1 | MT456405.1 | MK133097.1 |
| KM449067.1 | MH323175.1 | |

### LEMA DEMANGEI

MN845114.1

### LEMA PRAEUSTA

MN845117.1

### LEMA BORBONIAE

| | | |
|---|---|---|
| MT456402.1 | MT456404.1 | MT456403.1 |

### LEMA DIVERSIPES

| | |
|---|---|
| PQ540775.1 | PQ540776.1 |

### LEMA TRIVITTA

DQ001944.1

### LEMA YERBURYI

| | |
|---|---|
| PP667398.1 | PV467117.1 |

### LEMA SAIGONENSIS

| | |
|---|---|
| MN845116.1 | MN845115.1 |

### LEMA QUADRIPUNCTATA

PV056131.1

### LEMA IMMACULIPENNIS

DQ001939.1

### LEMA BITAENIATA

DQ001935.1

### LEMA HAMATA

DQ001938.1

### LEMA BIANNULARIS

DQ001931.1

### LEMA TRILINEA

DQ001945.1

### LEMA OBLITERATA
DQ001940.1

### LEMA FULVIPES
DQ001929.1

### LEMA BILINEATA
| | | |
|---|---|---|
| MT002928.1 | MT002929.1 | MT002941.1 |
| MT002930.1 | MT002931.1 | MT002943.1 |
| MT002932.1 | MT002934.1 | MT002945.1 |
| MT002935.1 | MT002936.1 | MT002933.1 |
| MT002937.1 | MT002946.1 | MT002944.1 |
| MT002942.1 | MT002947.1 | |

### LEMA FOVEIPENNIS
DQ001937.1

### LEMA BOUCHARDI
DQ001936.1

### LEMA REGULARIS
DQ001948.1

### LEMA INSULARIS
DQ001941.1

### LEMA SOLANI
| | | |
|---|---|---|
| Lema solani CO1 – PZ276999 | Lema solani 18S – PZ282238 | Lema solani 28S – PZ282240 |

### LEMA EQUESTRIS
| | | |
|---|---|---|
| Lema equestris CO1 – PZ276998 | Lema equestris 18S – PZ282237 | Lema equestris 28S – PZ282239 |

### OULEMA
| | | |
|---|---|---|
| KP763006.1 | KP763033.1 | DQ155800.1 |
| KP763086.1 | | |

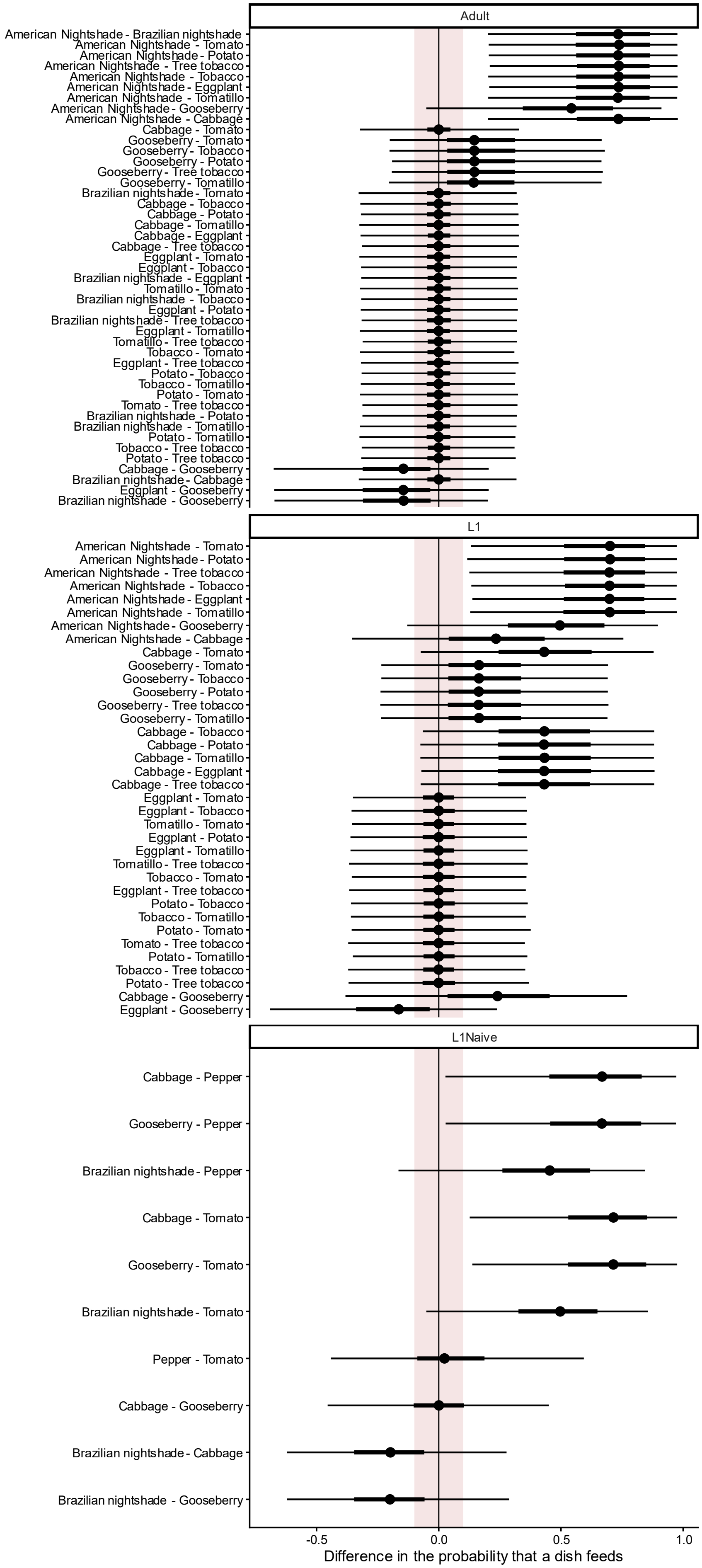
Adult
American Nightshade - Brazilian nightshade
American Nightshade - Tomato
American Nightshade - Potato
American Nightshade - Tree tobacco
American Nightshade - Tobacco
American Nightshade - Eggplant
American Nightshade - Tomatillo
American Nightshade - Gooseberry
American Nightshade - Cabbage
Cabbage - Tomato
Gooseberry - Tomato
Gooseberry - Tobacco
Gooseberry - Potato
Gooseberry - Tree tobacco
Gooseberry - Tomatillo
Brazilian nightshade - Tomato
Cabbage - Tobacco
Cabbage - Potato
Cabbage - Tomatillo
Cabbage - Eggplant
Cabbage - Tree tobacco
Eggplant - Tomato
Eggplant - Tobacco
Brazilian nightshade - Eggplant
Tomatillo - Tomato
Brazilian nightshade - Tobacco
Eggplant - Potato
Brazilian nightshade - Tree tobacco
Eggplant - Tomatillo
Tomatillo - Tree tobacco
Tobacco - Tomato
Eggplant - Tree tobacco
Potato - Tobacco
Tobacco - Tomatillo
Potato - Tomato
Tomato - Tree tobacco
Brazilian nightshade - Potato
Brazilian nightshade - Tomatillo
Potato - Tomatillo
Tobacco - Tree tobacco
Potato - Tree tobacco
Cabbage - Gooseberry
Brazilian nightshade - Cabbage
Eggplant - Gooseberry
Brazilian nightshade - Gooseberry
L1
American Nightshade - Tomato
American Nightshade - Potato
American Nightshade - Tree tobacco
American Nightshade - Tobacco
American Nightshade - Eggplant
American Nightshade - Tomatillo
American Nightshade - Gooseberry
American Nightshade - Cabbage
Cabbage - Tomato
Gooseberry - Tomato
Gooseberry - Tobacco
Gooseberry - Potato
Gooseberry - Tree tobacco
Gooseberry - Tomatillo
Cabbage - Tobacco
Cabbage - Potato
Cabbage - Tomatillo
Cabbage - Eggplant
Cabbage - Tree tobacco
Eggplant - Tomato
Eggplant - Tobacco
Tomatillo - Tomato
Eggplant - Potato
Eggplant - Tomatillo
Tomatillo - Tree tobacco
Tobacco - Tomato
Eggplant - Tree tobacco
Potato - Tobacco
Tobacco - Tomatillo
Potato - Tomato
Tomato - Tree tobacco
Potato - Tomatillo
Tobacco - Tree tobacco
Potato - Tree tobacco
Cabbage - Gooseberry
Eggplant - Gooseberry
L1Naive
Cabbage - Pepper
Gooseberry - Pepper
Brazilian nightshade - Pepper
Cabbage - Tomato
Gooseberry - Tomato
Brazilian nightshade - Tomato
Pepper - Tomato
Cabbage - Gooseberry
Brazilian nightshade - Cabbage
Brazilian nightshade - Gooseberry
-0.5
0.0
0.5
1.0
Difference in the probability that a dish feeds

Supplemental Figure 6. Pairwise contrasts in the probability that a dish feeds, within each life stage, for all plant pairs including those on which no feeding was observed. Points are posterior medians with 50% (thick) and 95% (thin) equal-tailed credible intervals; the red shaded band is the ROPE for a difference in probability (+/- 0.10).

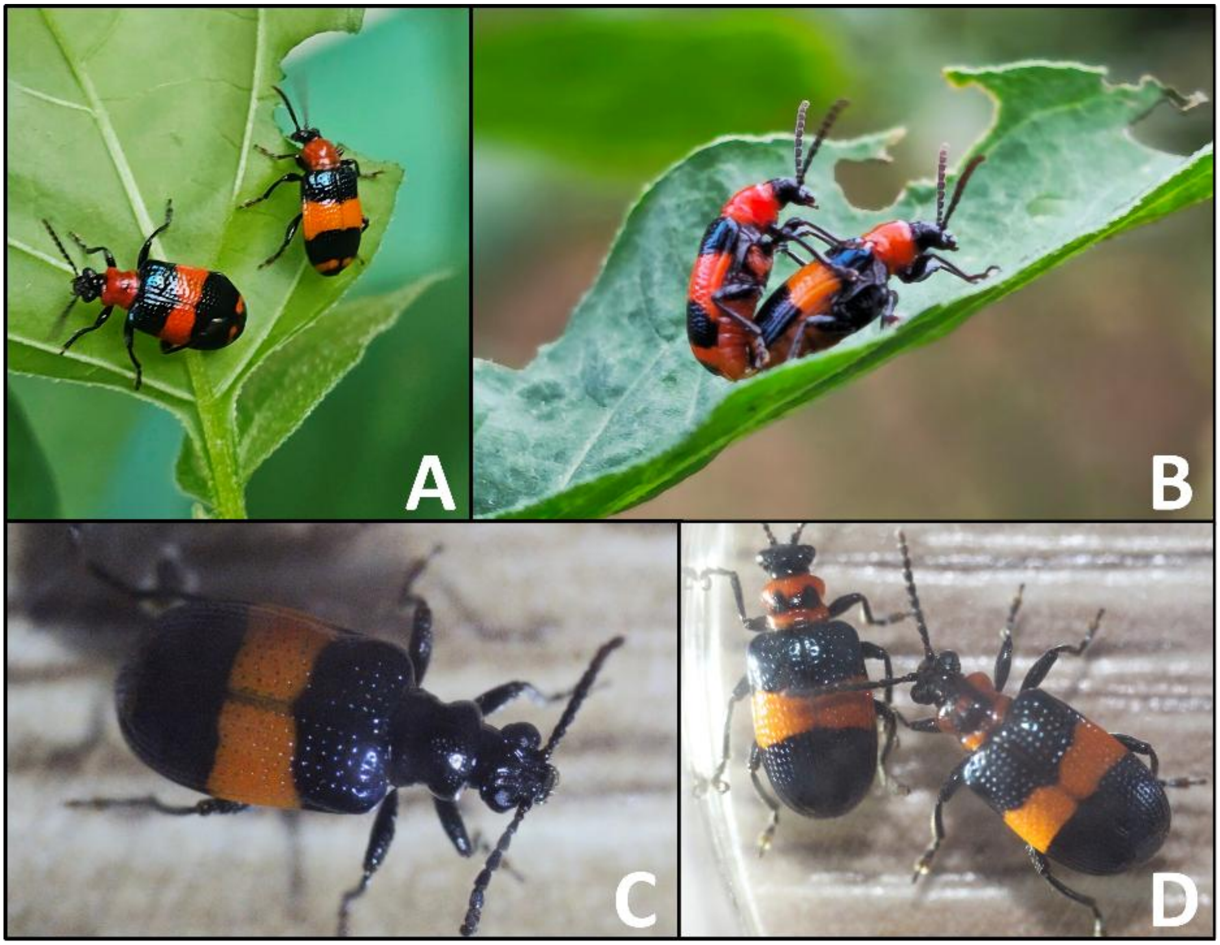


Supplemental Figure 7. A comparison of colour morphs of *L. equestris*. The top images show orange and rufous elytra phenotypes feeding on the same leaf (A) and mating (B). The thoracic collar varies in colour from red (A and B) to black (C), with some specimens showing intermediate colouration (D).

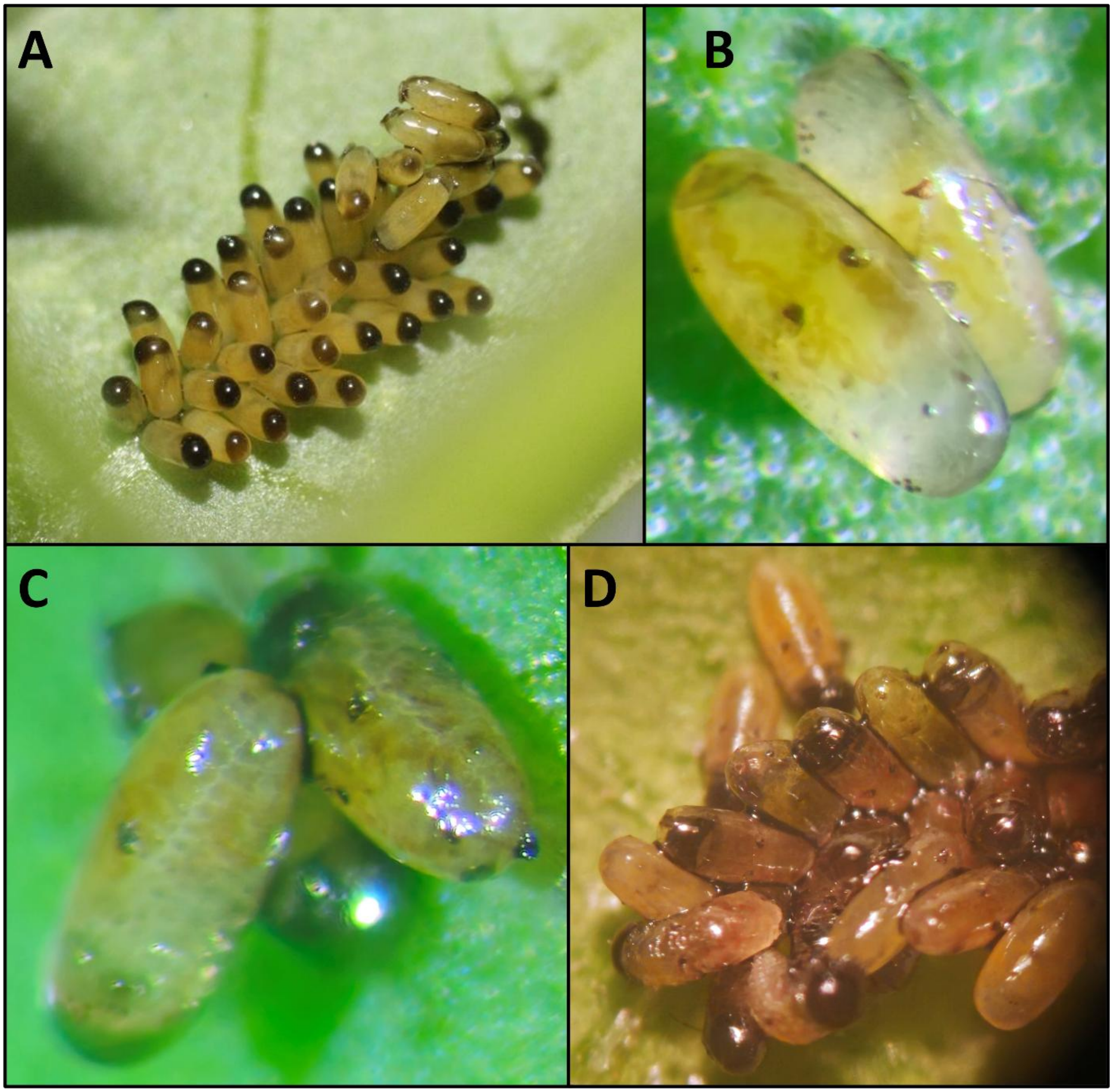


Supplemental Figure 8. Eggs of *L. equestris*. A. Newly deposited eggs along the midrib of the leaf. B. Eggs within 24 hours of hatching, with visible eyespots, abdominal spots, and malpighian tubules. C. Eggs within 12 hours of hatching that have begun to contract and have visible tracheae. D. A clutch of eggs comprised of emerging larvae and eggs, exhibiting sclerotized head capsules, that are within a couple of hours of hatching.

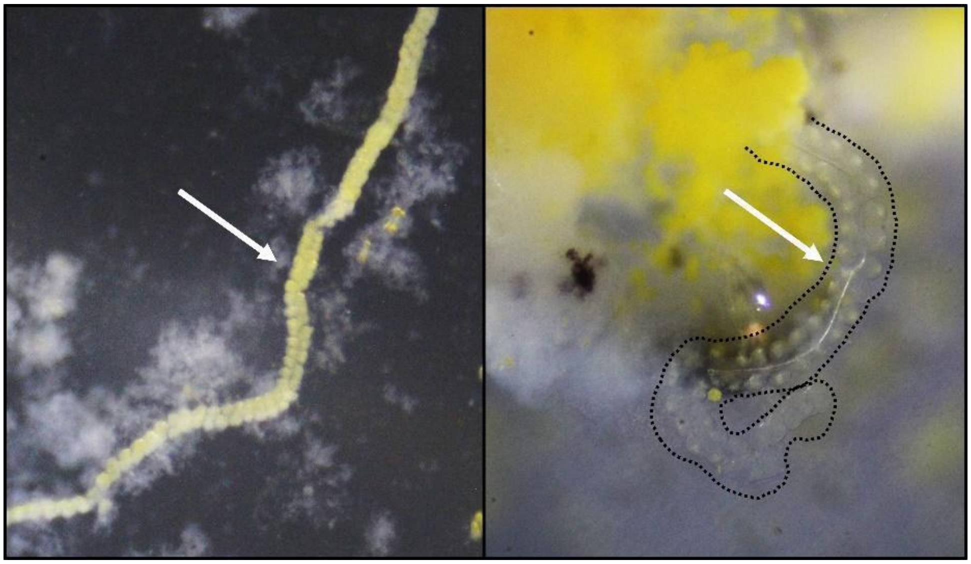

Supplemental Figure 9. Larval salivary glands from the feeding stage of an L4 larvae (left) and the prepupal stage (right).

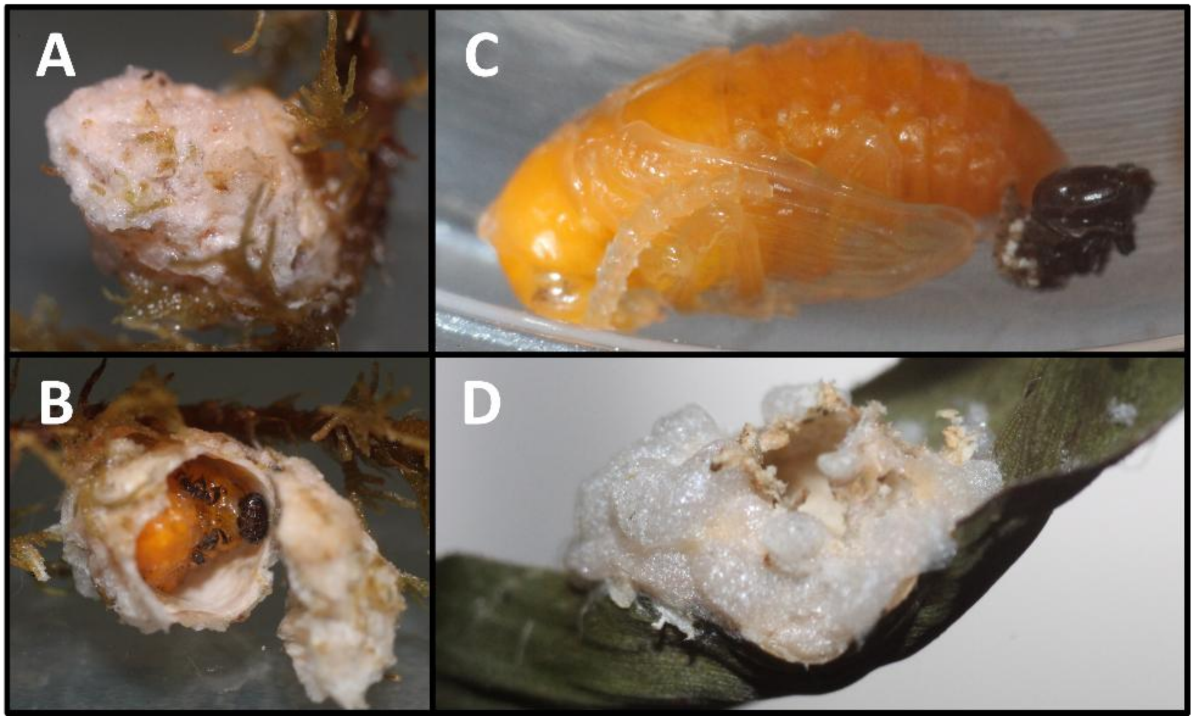


Supplemental Figure 10. Photographs of *Lema equestris* pupa and pupal cell. A. The foam constructed pupal cell. B. Prepupal larva in the pupal cell. C. Fully formed pupa. D. Pupal cell with the foam chewed open when the adult beetle emerged.

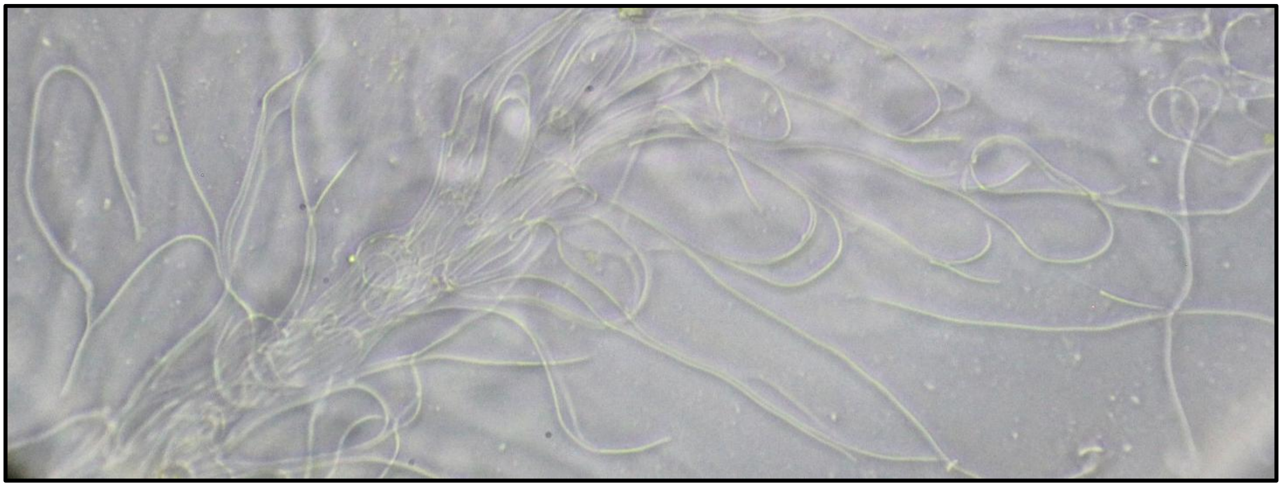

Supplemental Figure 11. Filamentous sperm of *L. equestris* viewed at 40x magnification.

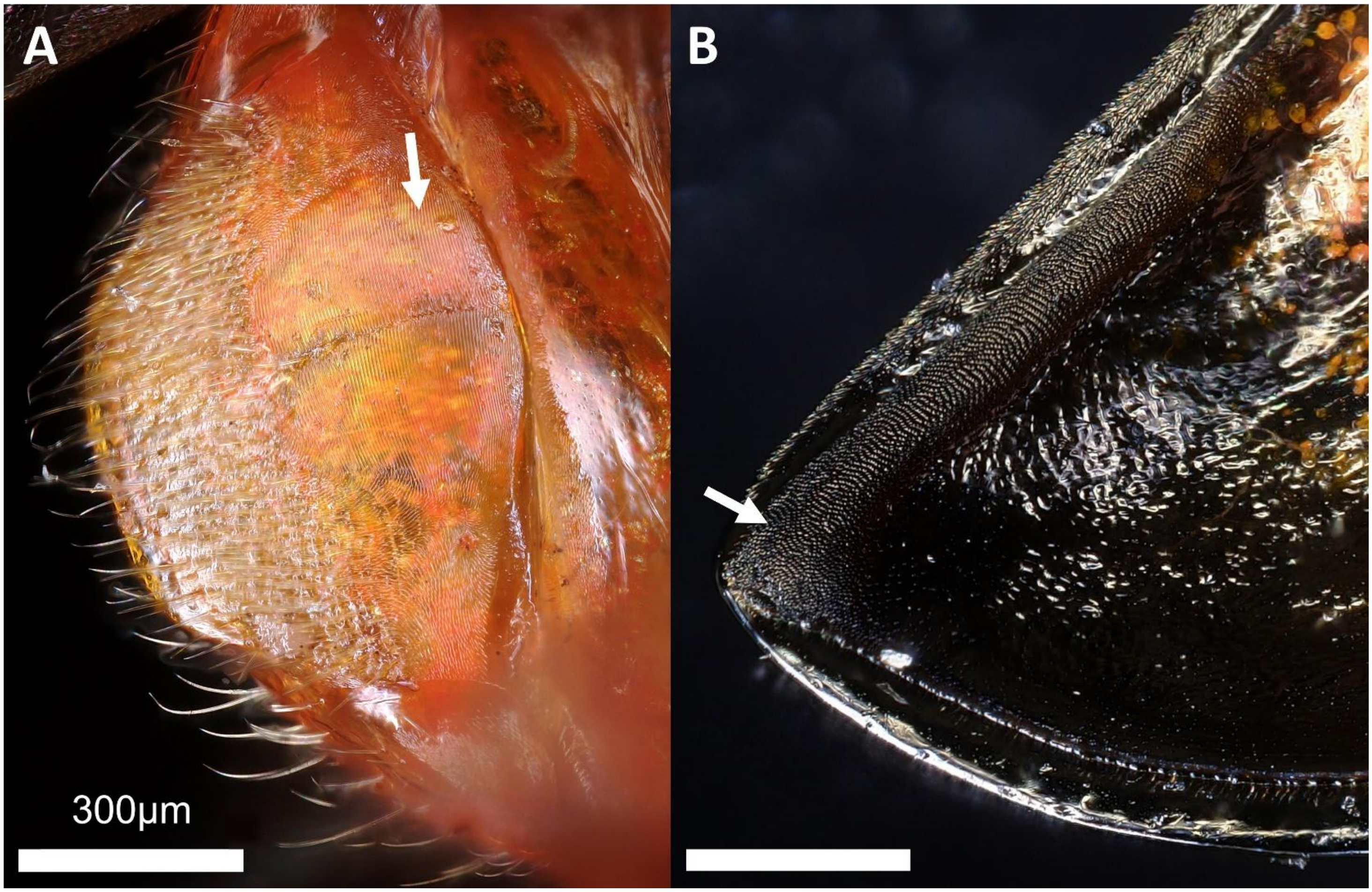


Supplemental Figure 12. Stridulatory rake and file of *L. equestris* with white arrows to highlight the structures. A. The dorsal rake on the basal portion of the posteriormost tergite. B. The stridulatory file at the distal end of the ventral side of the elytra.
